# The high turnover of ribosome-associated transcripts from *de novo* ORFs produces gene-like characteristics available for *de novo* gene emergence in wild yeast populations

**DOI:** 10.1101/329730

**Authors:** Éléonore Durand, Isabelle Gagnon-Arsenault, Johan Hallin, Isabelle Hatin, Alexandre K Dubé, Lou Nielly-Thibaut, Olivier Namy, Christian R Landry

## Abstract

Little is known about the rate of emergence of genes *de novo*, how they spread in populations and what their initial properties are. We examined wild yeast (*Saccharomyces paradoxus*) populations to characterize the diversity and turnover of intergenic ORFs over short evolutionary time-scales. With ~34,000 intergenic ORFs per individual genome for a total of ~64,000 orthogroups identified, we found *de novo* ORF formation to have a lower estimated turnover rate than gene duplication. Hundreds of intergenic ORFs show translation signatures similar to canonical genes. However, they have lower translation efficiency, which could reflect a mechanism to reduce their production cost or simply a lack of optimization. We experimentally confirmed the translation of many of these ORFs in laboratory conditions using a reporter assay. Translated intergenic ORFs tend to display low expression levels with sequence properties that generally are close to expectations based on intergenic sequences. However, some of the very recent translated intergenic ORFs, which appeared less than 110 Kya ago, already show gene-like characteristics, suggesting that the raw material for functional innovations could appear over short evolutionary time-scales.

## Introduction

The emergence of new genes is a driving force for phenotypic evolution. New genes may arise from pre-existing gene structures through genome rearrangements leading to gene duplication, gene fusion or horizontal gene transfer, or *de novo* from previously non-coding regions (Chen et al. 2013). *De novo* gene birth was considered highly unlikely (Jacob 1977) up until the last decade when comparative genomics approaches shed light on the role of intergenic regions as a regular source of new genes (Tautz and Domazet-Loso 2011; Landry et al. 2015; Schlotterer 2015; McLysaght and Hurst 2016). Compared to other mechanisms, *de novo* gene origination is a source of complete innovation because genes emerge solely from mutations, not from the modification of preexisting genes, with preexisting functions (McLysaght and Hurst 2016).

Non-coding regions need to go through three major steps to become gene-coding, the first two occurring in any order. i) The acquisition of an Open Reading Frame (ORF) by mutations conferring a gain of in-frame start and stop codons, and ii) the acquisition of regulatory sites to induce transcription and translation of the ORF. The third step is the retention of the expressed ORF and its selection because it encodes a less toxic or beneficial polypeptide (Schlotterer 2015; Nielly-Thibault and Landry 2018). The subsequent maintenance of the structure by purifying selection will lead to the gene being shared among species, as we see for groups of homologous canonical genes. The birth of genes *de novo* could in theory be a frequent process since numerous ORFs in non-annotated regions are associated with ribosomes, indicating that they are likely translated and thus have the potential to produce *de novo* polypeptides, which are the raw material necessary for *de novo* gene birth (Ingolia et al. 2009; Wilson and Masel 2011; Carvunis et al. 2012; Ruiz-Orera et al. 2014; Lu et al. 2017; Vakirlis et al. 2017; Ruiz-Orera et al. 2018). The different steps could be accelerated in some ways, depending on the genomic context. For instance, ORFs could emerge in long non-coding RNAs (lncRNAs) with relatively high pre-existing expression levels that reflect functions unrelated to the newly emerged ORF (Xie et al. 2012).

Many putative *de novo* genes have been identified (McLysaght and Hurst 2016), but there is generally limited information about their translation and only few have been functionally characterized (Begun et al. 2006; Levine et al. 2006; Begun et al. 2007; Cai et al. 2008; Zhou et al. 2008; Knowles and McLysaght 2009; Li et al. 2010; Baalsrud et al. 2017). These young genes are generally small with a simple intron-exon structure, they are on average less expressed than canonical genes and they may diverge rapidly compared to older genes (Wolf et al. 2009; Tautz and Domazet-Loso 2011). These properties make it challenging to differentiate *de novo* emerging genes from non-functional ORFs (McLysaght and Hurst 2016). The absence of sequence similarities of a given gene with known genes in other species is not sufficient evidence for *de novo* origination, since it could also be due to rapid divergence between orthologs. This confusion resulted in spurious *de novo* origin annotations, especially over longer evolutionary time-scale (Gubala et al. 2017). One way to overcome the problem is to identify *de novo* genes and the corresponding orthologous non-coding sequences in closely related populations or species through synteny, which gives access to mutations occurring during the gene birth process rather than long after the appearance of the *de novo* genes (Begun et al. 2006; Levine et al. 2006; Begun et al. 2007; Cai et al. 2008; Zhou et al. 2008; Knowles and McLysaght 2009; Li et al. 2010).

The process of *de novo* gene birth has been framed under various hypotheses that consider the role of selection as acting at different time points. The continuum hypothesis involves a gradual change in characteristics from non-genic to genic and was used to explain patterns related to sizes of intergenic ORFs (Carvunis et al. 2012). The preadaptation hypothesis predicts extreme levels of gene-like characteristics in young *de novo* genes, as was observed for intrinsic structural disorder (Wilson et al. 2017). The two models both depend i) on the distribution of properties (non gene-like versus gene-like) of random polypeptides within intergenic regions and ii) whether these properties correlate with the probability that the peptides will have an adaptive potential. Examining the distribution of properties of novel polypeptides early after their emergence – before they potentially lose their initial properties – is therefore important to determine which one of the two models could be supported.

Another question of interest is whether local composition along the genome can accelerate gene birth. The size of intergenic regions, their GC composition and the genomic context (e.g. spurious transcription) may affect the birth rate of *de novo* genes (Vakirlis et al. 2017; Nielly-Thibault and Landry 2018). A recent study on different yeast species found *de novo* genes to preferentially emerge in GC-rich genomic regions, in recombination hotspot and near divergent promoters (Vakirlis et al. 2017). Another feature that may affect emergence, but also loss, of *de novo* genes is mutation rate; differences in mutation rate would affect the overall turnover of *de novo* genes. Finally, because turnover itself may covary with sequence base composition, the properties of *de novo* genes could also be biased towards specific properties (Nielly-Thibault and Landry 2018).

Here we explore the contribution of intergenic genetic diversity in the emergence and retention of the raw material for *de novo* gene birth in wild *Saccharomyces paradoxus* populations. We focus on this yeast species because of its compact genome and close relatedness with the model species *Saccharomyces cerevisiae*. One advantage of *S. paradoxus* over *S. cerevisiae* is that the divergence of populations or lineages within species reflects natural events and not human domestication and human caused admixture since it has not been domesticated (Charron et al. 2014; Leducq et al. 2016). Most importantly, *S. paradoxus* harbors clearly defined lineages whose divergence times can be established and offers different levels of divergence that allow us to investigate recently emerged *de novo* genes. Finally, the use of natural populations may eventually allow for the connection between the evolution of *de novo* genes and key evolutionary processes such as adaptation and speciation, which have been intensively studied in *S. paradoxus* over the past few years (Charron et al. 2014; Naranjo et al. 2015; Leducq et al. 2016; Eberlein et al. 2017; Leducq et al. 2017; Weiss et al. 2018).

Using this model, we characterized the repertoire and turnover of ORFs located in intergenic regions (named hereafter iORFs), as well as the associated putative *de novo* polypeptides using ribosome profiling, and examined how the properties of putative polypeptides covary with their age and expression, and how they compare with those of canonical genes.

## Results

### A large number of intergenic ORFs segregate in wild *S. paradoxus* populations

We first investigated the diversity and turnover of ORFs located in intergenic regions, which we named iORFs, and their characteristics in wild *S. paradoxus* strains (Supplementary information). Because eukaryotic genomes are pervasively transcribed (David et al. 2006; Pelechano et al. 2013), and lncRNAs may produce peptides (Ruiz-Orera et al. 2014), we initially assumed that any iORF could have the ability to be translated and thus, could contribute to the process of *de novo* gene birth. We used 24 *S. paradoxus* strains that are structured in three main lineages named *SpA*, *SpB* and *SpC* (Charron et al. 2014; Leducq et al. 2016) and two *S. cerevisiae* strains as outgroups (Fig. 1, see Fig. S1 for strain names). These lineages cover different levels of nucleotide divergence, ranging from ~ 13 % between *S. cerevisiae* and *S. paradoxus* to ~2.27 % between the *SpB* and *SpC* lineages (Kellis et al. 2003; Leducq et al. 2016).

**Figure 1.**
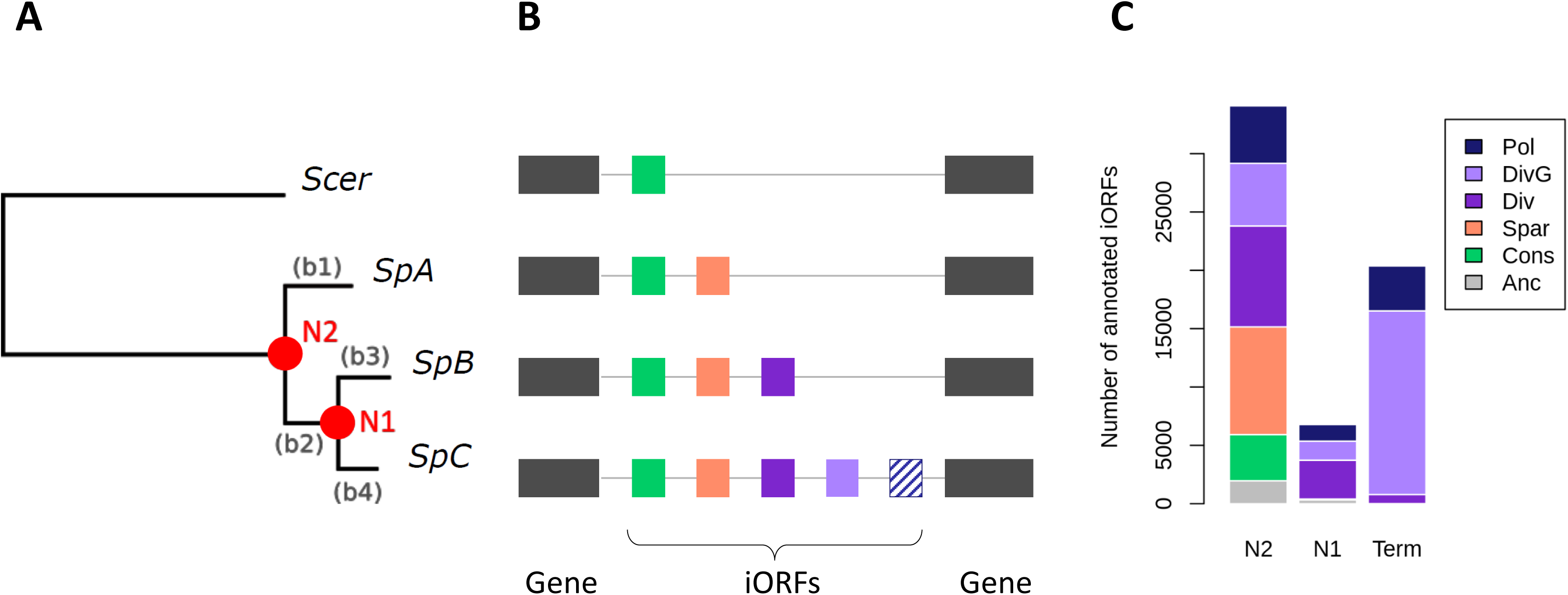
A large pool of iORFs segregate within and among *S. paradoxus* lineages. **A)** Phylogenetic tree of strains used for the reconstruction of ancestral intergenic sequences. Node and branch names are indicated in orange and grey respectively. **B)** Scheme of the iORFs annotation procedure (see Methods and Figure S1 for a complete description). Pairs of genes annotated as syntenic were used to align intergenic genomic regions in which iORFs were characterized. **C)** Number of annotated iORFs per age group, corresponding to the oldest node in which they were detected. ‘Term’ refers to iORFs appearing on terminal branches and being absent in ancestral reconstructions. iORFs are colored according to their conservation group (see Methods and Fig. S1): conserved (cons), S. paradoxus (Spar) specific and fixed, divergent (Div), divergent group-specific (DivG) and polymorphic (Pol). iORFs detected only in ancestral sequences are shown in gray.

We annotated iORFs as any first start and stop codons in the same reading frame not overlapping with known features, and with no minimum size (Carvunis et al. 2012; Sieber et al. 2018). We then measured the conservation of iORFs between strains using a conservative approach (Fig. S1, see Methods and Supplementary information). To understand how the iORF repertoire changes over a short evolutionary time scale, we also estimated the age of iORFs and their turnover using ancestral sequence reconstruction (see Methods).

We identified between 34,216 and 34,503 iORFs per *S. paradoxus* strain, for a total of 64,225 orthogroups annotated in at least one strain (Table 1 and Supplemental Table S1). We observed that the iORF repertoire of yeast populations is the result of frequent gains, losses, and size changes (Supplementary information). 56 % of the most ancient iORFs (detected at N2, Fig 1) are still segregating within *S. paradoxus*, showing the role of wild populations as a reservoir of iORFs that can be used to address the dynamics of early *de novo* gene evolution.

**Table 1.**
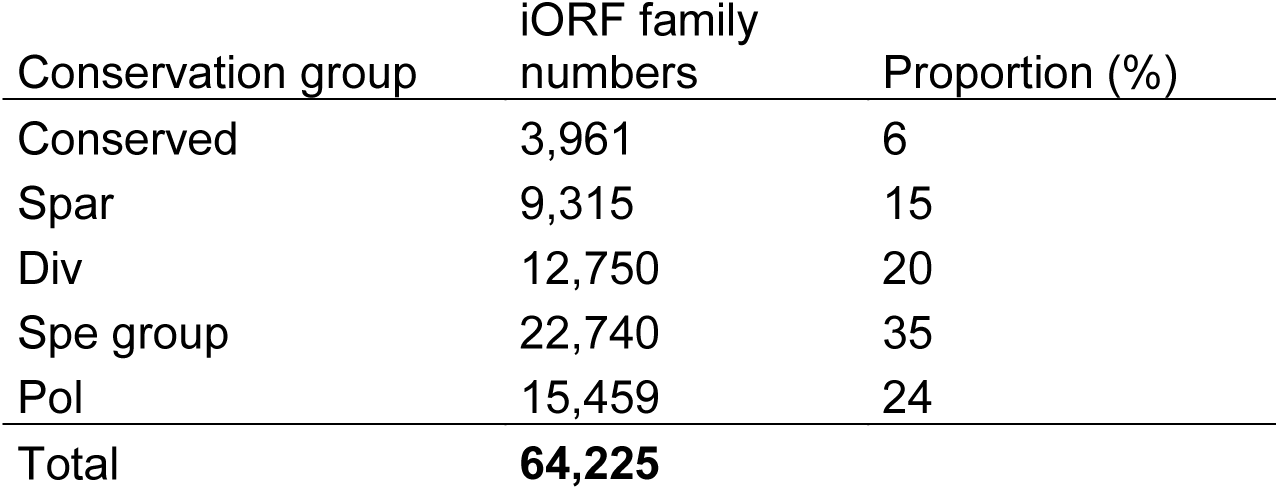
Number of iORFs per conservation group

### Hundreds of intergenic ORFs show signatures of active translation

We performed ribosome profiling to identify iORFs that are potentially translated and that thus possibly produce polypeptides. Only iORFs with a minimum size of 60 bp were considered for this analysis. Among them, 12 that displayed a significant blast hit when searched in the proteomes of 417 species, including 237 fungi, were removed for the downstream analysis (see Methods). The final set examined consisted of 19,689 iORFs. We prepared ribosome profiling sequencing libraries for four strains, one belonging to each lineage or species: YPS128 (*S. cerevisiae*), YPS744 (*SpA*), MSH-604 (*SpB*) and MSH-587-1 (*SpC*), in two biological replicates. All strains were grown in synthetic oak exudate (SOE) medium (Murphy et al. 2006) to approximate the natural conditions in which *de novo* genes could emerge in wild yeast strains.

Typically, a ribosome profiling density pattern is characterized by a strong initiation peak located at the start codon followed by a trinucleotide periodicity at each codon of protein-coding ORFs. We used this feature to identify a set of iORFs that are the most likely to be translated and we compared their translation intensity with that of annotated genes. We first detected peaks of initiation sites around start codons. As expected, the number of ribosome profiling reads located at this position is on average lower for iORFs than for annotated genes (Fig. 2A). However, we observed an overlap between the two read density distributions, illustrating a similar read density between highly expressed iORFs and lowly expressed genes. We observed an initiation peak for 73.9 to 87.9 % of standard annotated genes depending on the strain, and for 1.4 to 6.9 % of iORFs (Table 3 and Fig. 2B). Detected peaks were classified using three levels of precision and intensity: ‘p1’ for less precise peaks (+/-1nt relative to the first base of the start codon), ‘p2’ for precise peaks (detected at the exact first base of the start codon) and ‘p3’ for precise peaks with strong initiation signals characterized here by the highest read density in the ORF (see Methods). Among all iORFs with a detected initiation peak, 30, 35 and 34% respectively belong to p1, p2 and p3. A comparable repartition (Chi-square test, p-value= 0.59) was observed for annotated genes with 24, 40 and 36% for each precision group, showing that the precision levels used in our analysis are reliable.

**Figure 2.**
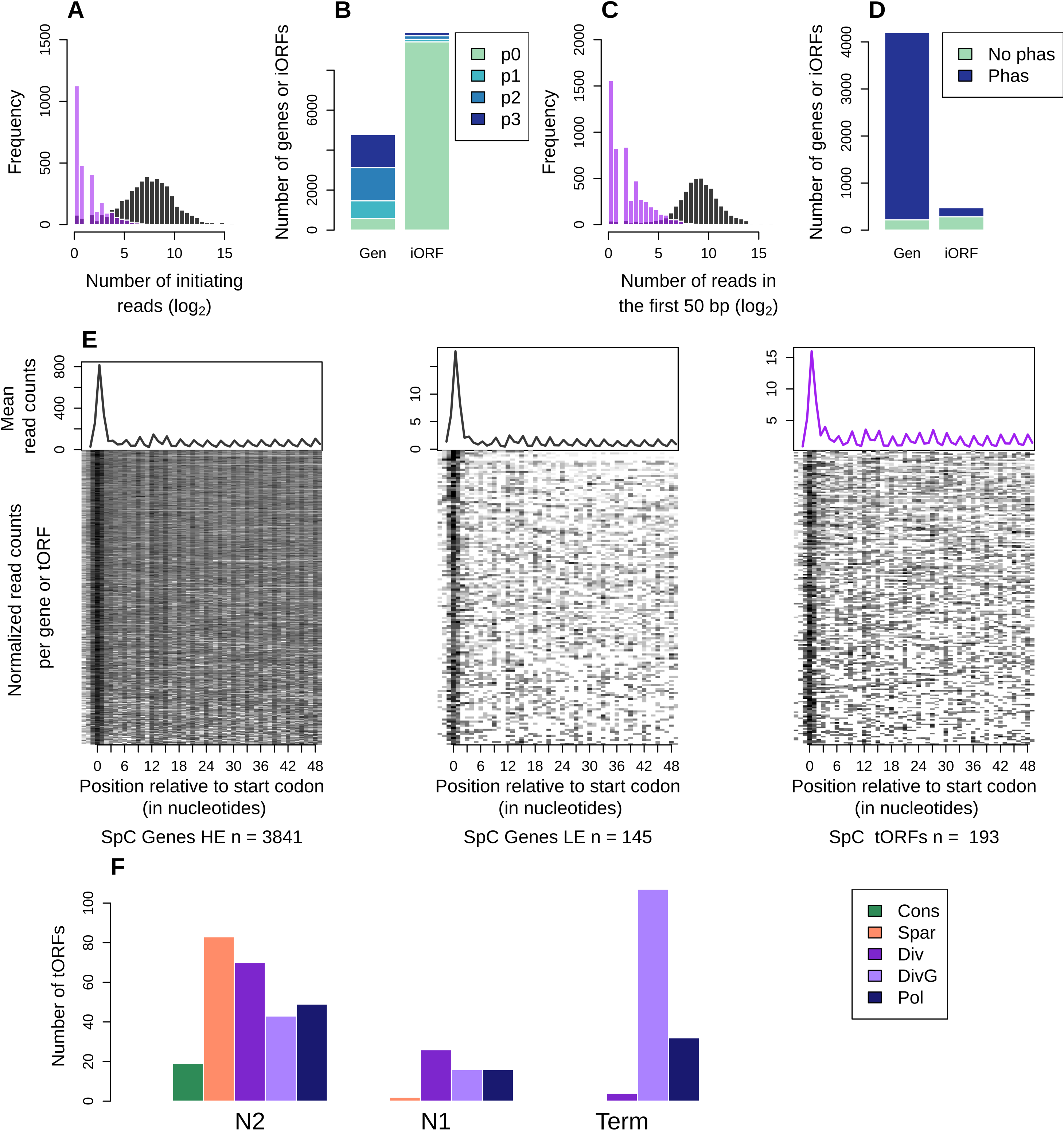
A fraction of the iORFs display translation signatures similar to genes. **A)** Distribution of the ribosome profiling read counts for genes (grey) and iORFs (purple) at the start codon position. **B)** Number of genes (Gen) or iORFs with a detected initiation peak at the start codon position. Peaks are colored according to the precision of the detection (see Methods), from the most precise (p3) to the least precise (p1). Genes and iORFs with no peaks detected are shown in green (p0). **C)** Distribution of the ribosome profiling read counts in the first 50 nt of iORFs excluding the start codon **D**) Proportions of genes or iORFs with a significant in frame codon periodicity (read phasing in blue) among genes and iORFs with a detected initiation peak. Genes and iORFs with no detected phasing are shown in green. **E)** Metagene analysis for significantly translated highly (HE, left) or lowly (LE, middle) expressed genes (grey), and intergenic translated ORFs (tORFs) (purple, right). The mean of the 5’ read counts is plotted along the position relative to the start codon for significantly translated genes or tORFs. The lines of the matrix indicate the normalized coverage of genes or tORFs with significant translation signatures, with one feature per line. **A-E**) Results for the *SpC* strain MSH-587-1 are shown (see Fig. S2 for *SpA* and *SpB* results). **F)** Total Number of tORFs per conservation group per age detected in all sequenced strains.

We measured codon periodicity, which is characterized by an enrichment of reads at the first nucleotide of each codon in the first 50 nt excluding the start codon. As for the start codon region, the number of ribosome profiling reads is lower for iORFs compared to known genes (Fig. 2C). Among the features with a detected initiation peak, 91.8 to 94.8% of genes and 29.4 to 41 % of iORFs show a significant codon periodicity per strain (Table 3 and Fig. 2D). The number of detected translation signals is lower in the *SpB* strain, which is most likely due to a lower number of reads obtained for this strain and the use of raw read density in this analysis (see Methods). iORFs with an initiation peak and a significant periodicity in at least one strain were considered as significantly translated and labeled tORFs, whereas iORFs with no significant translation signatures were labeled ntORFs. We performed a metagene analysis on annotated genes and tORFs, which revealed a similar ribosome profiling read density pattern between low expressed genes and tORFs, and confirmed a distinct codon periodicity with significant translation signature for tORFs (Fig. 2E and S2). The resulting tORF set contains 418 orthogroups with sizes ranging from 60 to 369 nucleotides. They are represented in all age and conservation categories, which suggests a continuous emergence of potentially translated ORFs along the phylogeny (Fig. 2F).

We compared our results with those resulting from an alternative method (RiboTaper) that is based on the quantification of the in-frame nucleotide periodicity to detect *de novo* translated ORFs (Calviello et al. 2016). Among the 418 tORFs detected in our primary analysis, 170 were also annotated *de novo* with RiboTaper (Fig. S3). Additionally, we detected 373 translated ORFs private to the RiboTaper method. Compared with tORFs private to our methods, tORFs private to RiboTaper are also characterized by an overall clear trinucleotidic periodicity but they display on average weaker initiation peaks as well as an overall lower ribosome profiling read coverage in the first 50 bp (Fig. S3). Below, we describe the results of the analysis performed with the set of 418 tORFs detected using our method, and which have been confirmed using the subset of 170 tORFs detected by both methods, as well as with the set of 525 tORFs detected using RiboTaper (see Supplementary information and Supplemental Table S2). tORFs represent a small fraction (~2 %) of the 19,689 iORF orthogroups longer than 60 nt. This percentage may be a conservative estimate because the detection depends on the chosen method the conditions examined, the filters and the ribosome profiling sequencing depth. However, the number of tORFs is consistent with pervasive transcription measurements in *S. cerevisiae*, with several hundreds of transcripts detected in non-annotated genomic regions (David et al. 2006). Overall, for a genome of about 5,000 genes, the roughly 400 *de novo* tORFs which may produce *de novo* polypeptides could be an important contribution to the proteome diversity of these natural populations.

### Translational buffering acts on intergenic ORFs

We compared the expression levels of ancient and recent tORFs with that of known genes to examine if *de novo* polypeptides display gene-like expression levels. Because the *de novo* gene birth process under the continuum hypothesis involves an increase of tORF size, we also compared tORF properties while controlling for size ranges per age. The overlap between the size distributions of tORFs and genes is at the extremes of both distributions and the number of tORFs is not large enough to generalize the overall properties of longer tORFs with those of smaller genes (Fig. 3A).

**Figure 3.**
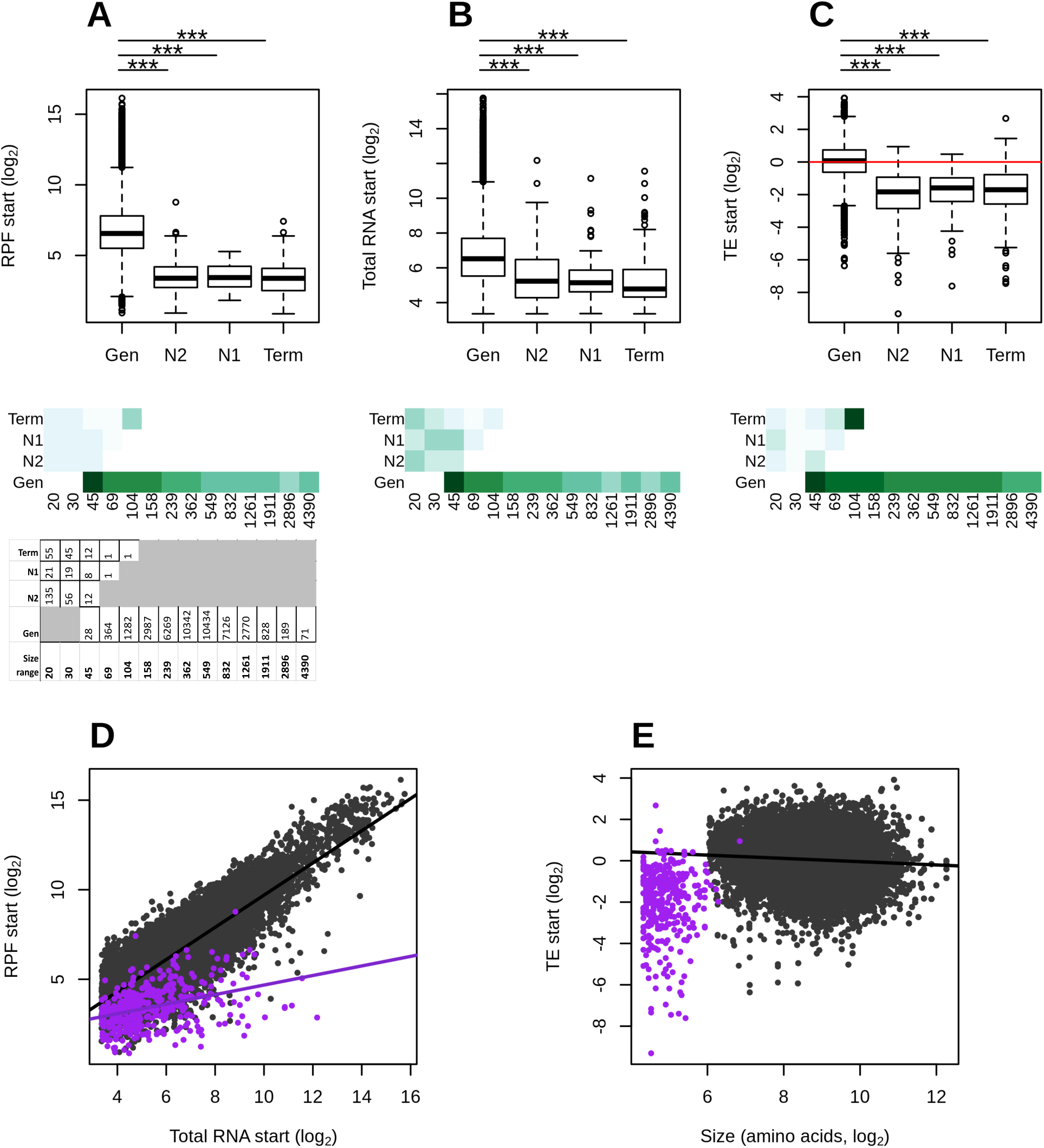
Putative intergenic polypeptides are less efficiently translated compared to genes. **A-C)** Ribosome profiling (RPF), total RNA and translation efficiency (TE) - read counts in the first 60 nt, normalized to correct for library size differences in log_2_ - are displayed for genes (Gen) and tORFs depending on their ages (N2, N1 and Term). Significant differences in pairwise comparisons are displayed above each plot (Wilcoxon test, *** for p-values < 0.001, ** for p-values < 0.01 and * for p-values < 0.05). Mean estimates per size range are colored by green intensities (from pale for low values to dark green for high values) below. Numbers per size range and age are indicated below the graph. **D)** RPF plotted as a function of total RNA for tORFs in purple, or genes in grey **. E)** TE plotted as a function of tORF or gene sizes (number of amino acid residues in log2). Regression lines are plotted for significant Spearman correlations (p-values < 0.05). Expression levels were calculated using the mean of the two replicates.

We investigated translation and transcription levels using ribosome profiling and total RNA sequencing. We estimated translation efficiency (TE) per gene and tORF as the ratio of ribosome profiling footprints (RPFs) over total mRNA normalized read counts. TE values increase with the number of translating ribosomes per molecule of mRNA, illustrating a more effective translation per mRNA unit (Ingolia et al. 2009). Note that RPF and total RNA coverages were calculated on the first 60 nt for both genes and tORFs to reduce the bias introduced by the high number of reads at the initiation codon compared to the rest of the sequences, which tends to increase TE estimates in short tORFs compared to longer genes. After this correction, TE values remain significantly correlated with gene size but the effect is small and should not interfere in our analysis (Fig. 3E).

As expected for intergenic regions, tORFs were less transcribed and translated than genes (Wilcoxon test, p-values < 2.2x10^-16^, Fig. 3A-B). We also observed, on average, a significantly lower TE (Wilcoxon test, p-value = < 2.2x10^-16^, Fig. 3C) for most of tORFs compared to genes, suggesting that tORFs are less actively translated than genes, even when considering the same size ranges (Fig. 3C). We note, however, that the longest tORF size range category contains only one tORF (tORF_102655), which displays a much higher TE value compared with tORFs from all other size ranges.

More generally, the most highly transcribed tORFs display a more reduced TE compared to genes (Fig. 3D, ANCOVA, p-value < 2.2×10^-16^). This buffering effect was confirmed when considering the 525 tORFs detected using RiboTaper, as well as in the subset of 170 tORFs detected using both methods (Fig. S4). A potential consequence of this post-transcriptional buffering is a reduction of polypeptides translated per molecule of mRNA. The buffering of highly transcribed tORFs may be due to a rapid selection to reduce the production of toxic polypeptides or may simply be a consequence of a recent increase in transcription without a change in features that would increase translation rate (e.g. codon usage). The buffering effect is similar among tORFs of different ages, with no significant pairwise differences between slopes (data not shown), which supports the hypothesis of no selection for or against translation. Although, on average, tORFs have lower expression levels and TE values than genes, we noted a significant overlap between expression levels and TEs in the two sets, which means that some tORFs have gene-like expression levels and translation efficiencies.

### Translated intergenic polypeptides display a high variability for gene-like traits

A recent study suggested that selection favors pre-adapted *de novo* young genes with high levels of intrinsic protein structural disorder (ISD). They showed that young *de novo* genes were more disordered than old genes, whereas random polypeptides in intergenic regions were on average less disordered (Wilson et al. 2017). This would suggest that young polypeptides with an adaptive potential would already be biased in terms of structural properties compared to the neutral expectations based on random sequences. We used our data on within species diversity to examine whether such features are indeed present among tORFs. We examined the properties of predicted polypeptides as a function of emergence timing in order to follow evolution before or at the early beginning of the action of selection. We compared the level of intrinsic disorder, GC-content and genetic diversity (based on SNPs density) in tORFs as a function of age and with the properties of annotated known genes. We noted that these properties were confirmed when considering the 525 tORFs detected using RiboTaper, as well as in the subset of 170 tORFs detected using both methods (Fig. S5). On average, protein disorder and GC-content are lower in tORFs than in canonical genes regardless of the tORFs age (Wilcoxon test, p-values < 0.001, Fig. 4B-C). This pattern was confirmed for most of tORFs and genes sharing the same size range of 45-100 amino acid long (Fig. 4B-C).

**Figure 4.**
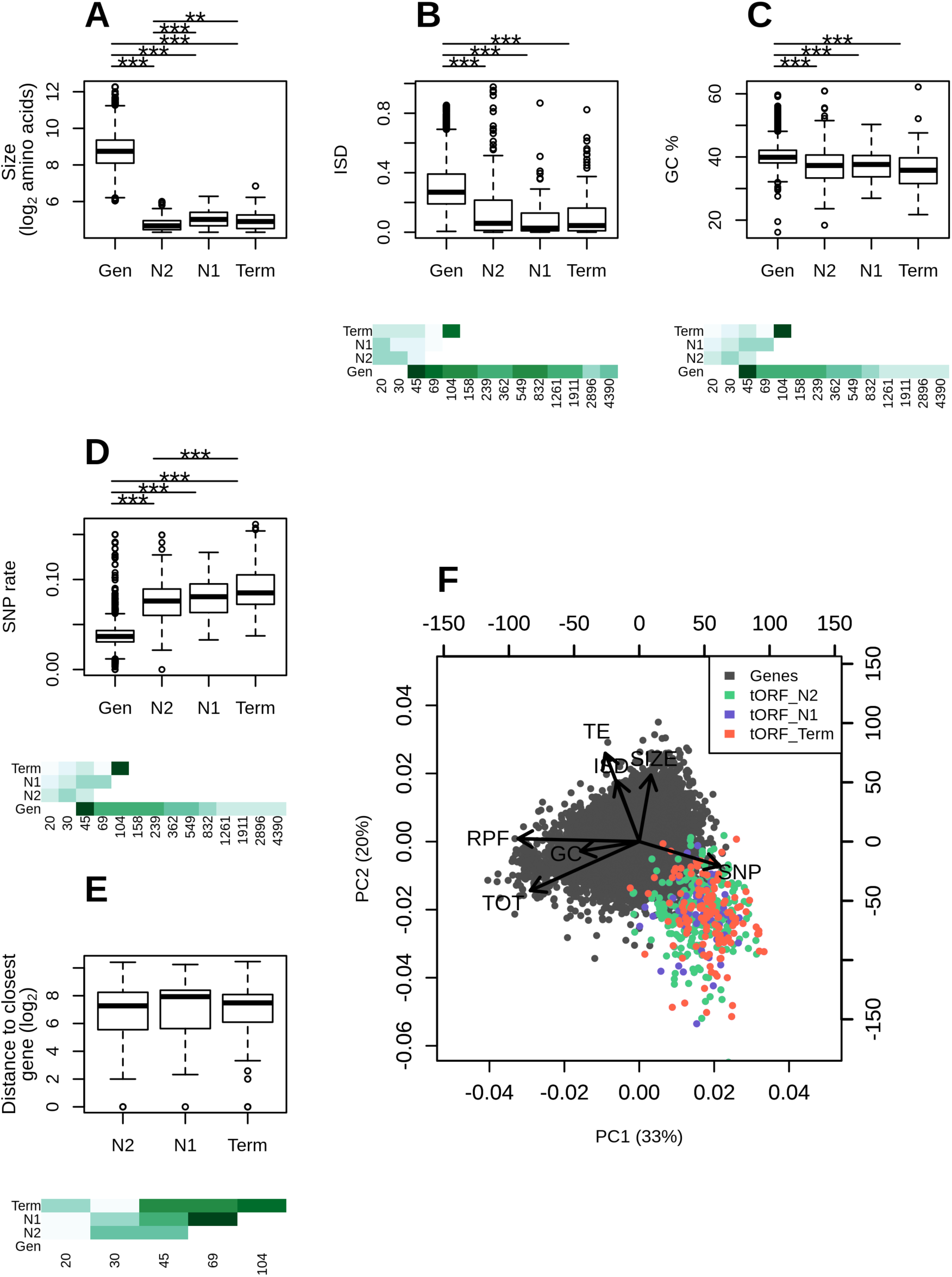
Age-dependent characteristics of intergenic polypeptides. **A-E)** Sizes (log_2_ number of residues), mean disorder (ISD), GC %, SNP density and distance to the closest gene are displayed for genes and tORFs as a function of their ages (N2, N1 and Term). Pairwise significant differences are displayed above each plot (Wilcoxon test, *** for p-values < 0.001, ** for p-values < 0.01 and * for p-values <0.05). Mean estimates per size ranges are colored with green intensities (from pale for low values to dark green high values) below. **F)** Principal component analysis using the number of residues (SIZE in log_2_), ribosome profiling (RPF), total RNA (TOT) and translation efficiency (TE) (as read counts in the first 60 nt normalized to correct for library size differences and in log_2_), intrinsic disorder (ISD), the GC% and SNP density (SNP). tORFs are colored as a function of their ages. The two first axis explain 33 and 20 % of the variation (total 53 %).

We examined if SNP density variation along the genome could influence tORF turnover. Regardless of their ages, tORFs are located in regions displaying a higher SNP density compared to genes, which is consistent with stronger purifying selection on canonical genes (Fig. 4D). Moreover, younger tORFs, appearing along the terminal branches, tend to be in regions with higher SNP rates compared to older ones at N2, even when considering the same size ranges (Fig. 4D). This may be due to mutation rate variation or differences in evolutionary constraints acting on tORF in an age specific manner. Older tORFs are not preferentially located at the proximity of genes where selection may be stronger (Fig. 4G), suggesting that the lower diversity observed at N2 is mainly due to a lower mutation rate. These observations suggest that younger tORFs are more likely to occur in rapidly evolving sequences with higher mutation rates. We performed a multivariate analysis to look for polypeptides with extreme values for multiple traits as an indicator of their functional potential. We observed a subset of tORFs sharing all characteristics that are typically considered to be gene-like in both more ancient or recent tORFs (Fig. 4F). Among them, tORF_102655, which is the only representative of the longest tORF size range on Fig. 3 and 4, is characterized by multiple gene-like characteristics with extreme intrinsic disorder, GC%, SNP rate and TE values (Fig. 3 and Fig. 4). This tORF, acquired along the *SpC* terminal branch and fixed in all strains of the *SpC* lineage, might be recruited by natural selection if gene-like characteristics increase its functional potential. Sequences are too similar between strains to test for purifying selection individually on each tORF. Instead, we estimated the likelihood of the global dN/dS ratio for two merged set of tORFs, containing ancient tORFs conserved in all *S. paradoxus* strains (set 1) or tORFs appearing at N1 and conserved between the *SpB* and *SpC* lineages (set 2). Both sets seem to evolve neutrally without significant purifying selection (NS p-values). Altogether, tORFs do not display significant purifying selection, but it appears that as a neutral pool, they provide raw material with gene-like characteristics for selection to act.

### Some intergenic translated ORFs display strong expression changes between lineages in SOE conditions

Our analysis has so far revealed that natural populations are provided with a regular supply of *de novo* putative polypeptides in intergenic regions (Table 2) at a rate sufficient to provide lineages that diverged less than 500,000 years ago with different gene contents. We looked for lineage-specific emerging putative polypeptides among tORFs based on significant differences of ribosome profiling coverage between each pair of strains (see Methods). Note that a translation gain or increase may be due to an iORF gain, a transcription/translation increase, or both. 33 tORFs display a significant lineage-specific expression increase, with 20, 5 and 8 tORFs in *SpA*, *SpB* and *SpC* respectively (Fig. 5 and S6). Among them, 24 are lineage-specific, and 16 of those were acquired along terminal branches, like the *SpB*-specific tORF_70680 (Fig. 5). Nearly 70 % of strong lineage-specific expression patterns are correlated with the presence of the tORF in one lineage only. This suggests that iORF turnover (gain and loss of start and stop codons) mostly explain translation differences and not a lineage expression increase in a region already containing a conserved iORF for instance. Three tORFs are more expressed in both *SpB* and *SpC* strains compared to *SpA* and *Scer*, suggesting an event occurring along branch b2 (Fig. 1A and S6). We also detected older expression gain/increase events in *S. paradoxus* relative to *S. cerevisiae* for 9 tORFs, for instance tORF_69174 (Fig. 5 and S6).

**Table 2.** Estimated age of iORFs in *S. paradoxus* lineages

| Age (Node or branch)<sup>1</sup> | Total | Numbers > or equal to 60nt<sup>2</sup> | Numbers with translation signature<sup>2</sup> |
| --- | --- | --- | --- |
| N2 | 34,092 | 8,336 | 221 |
| N1 | 6,782 | 2,664 | 56 |
| b1 (*SpA*) | 8,454 | 3,608 | 73 |
| b3 (*SpB*) | 6,860 | 2,948 | 13 |
| b4 (*SpC*) | 5,324 | 2,235 | 48 |
| Total without redundancy<sup>2</sup> | 61,243 | 19,689 | 418 |

<sup>1</sup> N1 and N2 refer to phylogenetic nodes (see Fig. 1A). b1, b3 and b4 are terminal branches, these categories refer to iORFs absent in ancestral sequences (based on the conservation of the start and stop position in the same reading frame). iORFs present in none of the strains used for reconstruction analysis were removed (see Methods).
<sup>2</sup> The 12 iORFs with significant blastp hits against reference proteomes (see results and Methods) were removed.

**Table 3.**
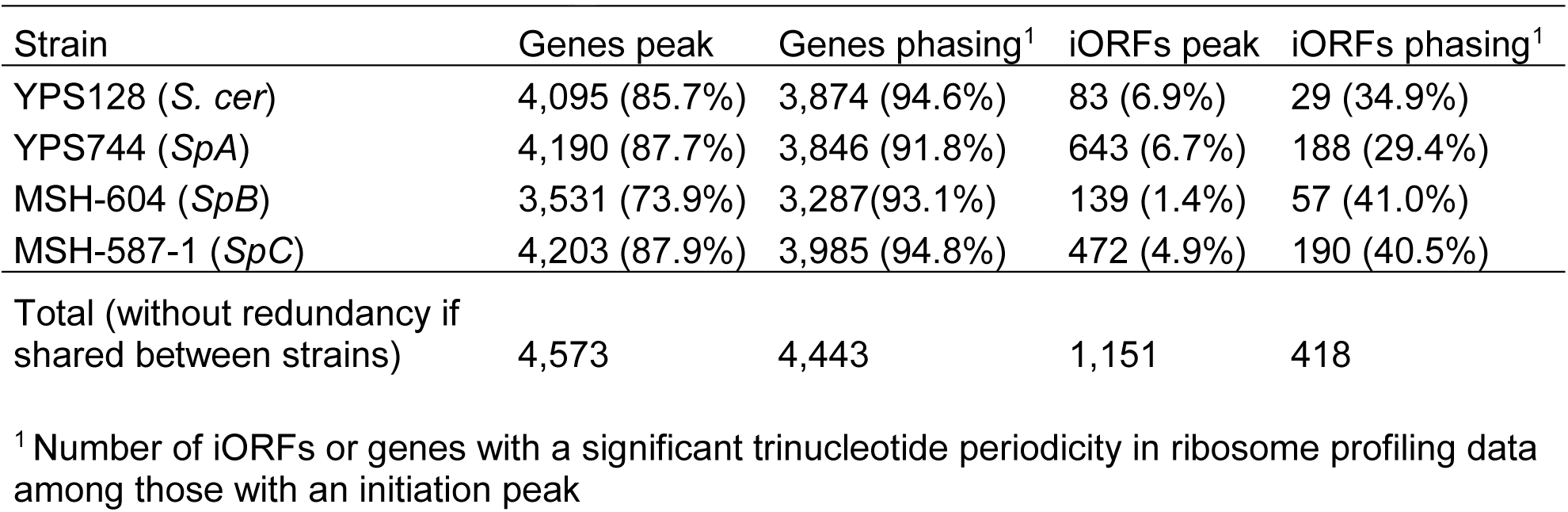
Detection of translated genes or iORFs

**Figure 5.**
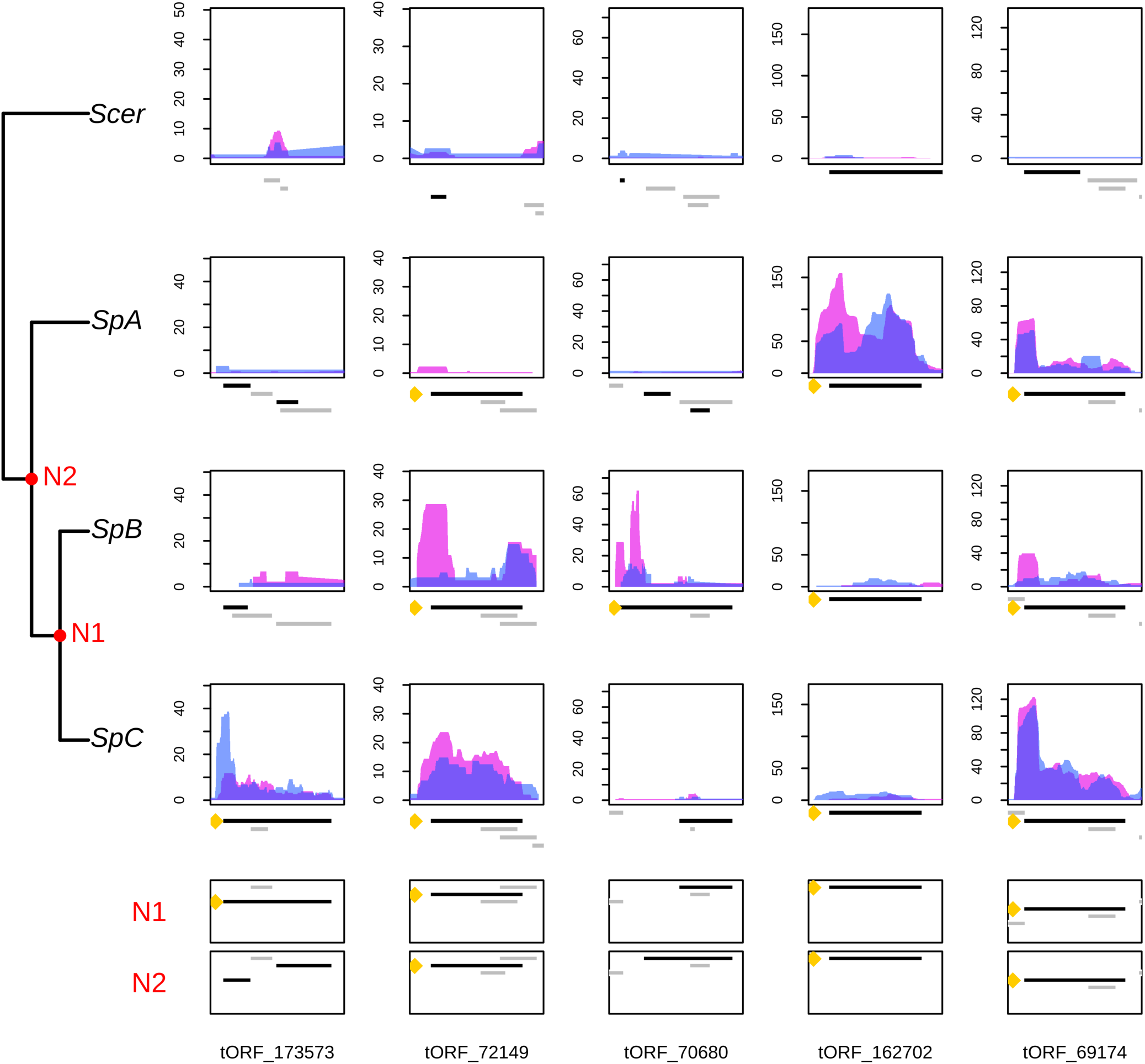
A continuous emergence of putative polypeptides in *S. paradoxus*. Normalized RPF read coverage for a selection of lineage specific (or group specific) tORFs per strain. RPF read coverages are displayed for replicate 1 and 2 with a blue or pink area respectively. The positions of all iORFs (including ntORFs and tORFs) in the genomic area are drawn below each plot. The tORF of interest is labeled with a yellow dot and is plotted in black. iORFs overlapping the iORF of interest are plotted in black when they are in the same reading frame, and in grey when they are in a different reading frame as the selected tORF.

### Several tORFs show significant translation using a reporter assay

Finally, we selected the 45 tORFs displaying significant translation changes described above to test for translation using a reporter gene. We chose to cover ancient and recent polypeptide gain events (i.e. lineage-specific or older events). We used a mutated dihydrofolate reductase gene (DHFR) as a reporter enzyme to fuse with the tORFs (Tarassov et al. 2008; Freschi et al. 2013). This enzyme confers resistance to methotrexate (MTX) when expressed at significant levels. We integrated the DHFR coding sequence that excludes the start codon in fusion at the 3’ end of the candidate tORFs in *SpA*, *SpB* and *SpC* genetic backgrounds. We fused the DHFR in the same reading frame as the tORF (construction tORF_DHFR_in_frame) to test for translation controlled by the native tORF promoter and most likely translation initiation codon (Fig. 6). We also fused the DHFR with the tORFs in a different reading frame as a negative control (tORF_DHFR_out_of_frame). We then tested the translation of the constructs using cell growth assay on a medium supplemented with MTX and on a medium supplemented with DMSO as a control (Fig. 6) (Methods). We also fused the DHFR with 12 canonical genes as positive controls.

**Figure 6.**
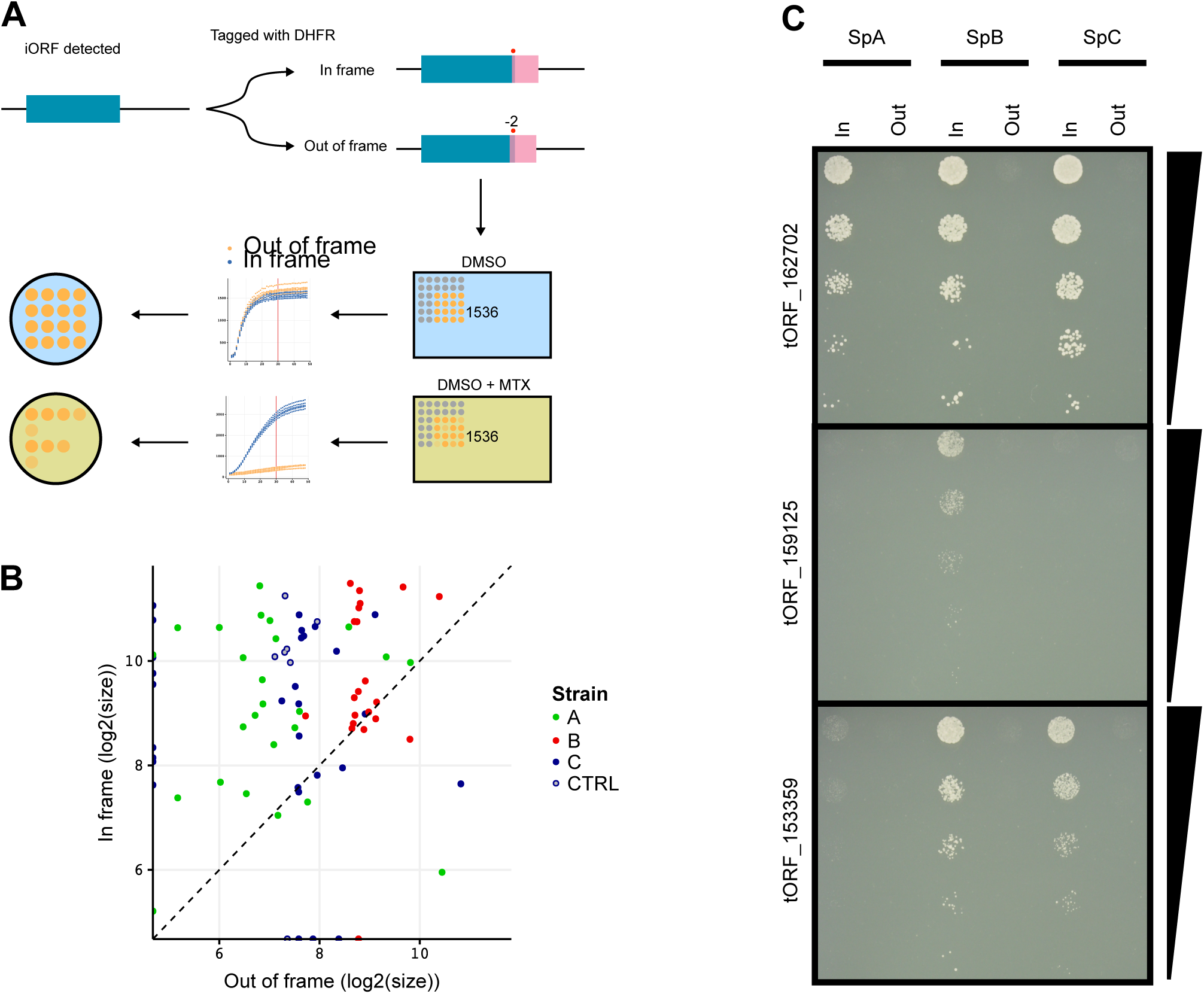
DHFR tagging confirms expression of tORFs. **A)** Conceptual figure of the approach, 45 tORFs were tagged with a full-length DHFR, in frame of out of frame in *SpA*, *SpB* and *SpC*, then phenotyped by time-resolved imaging and spot-dilution assays. **B)** Log_2_ colony sizes of strains tagged with DHFR in frame (y-axis) and out of frame (x-axis). The colony size is taken after 60 hours of growth (shown as a red vertical line in panel A) on medium supplemented with methotrexate. Colors represent the different strains, the CTRL strains are tagged in canonical genes, these constructs were made in the *SpC* strain. Dashed line: y = x. **C)** Spot-dilution assays further confirm expression of the tORFs, and shows differential expression of tORF_159125 and tORF_153359. 10-fold dilutions go from top to bottom. **B, C)** For the corresponding controls in medium not supplemented with methotrexate, see Fig. S7.

We found support for the translation of 26 of the 45 tORFs in at least one strain (Fig 6 and Fig. S8) and 6 tORFs with a translation signal potentially from a different reading frame, where out of frame fusion cells grew better on selective medium than in frame fusions (Fig. 6, Fig. S7 and Fig. S8). Interestingly, four of these tORFs have overlapping iORFs in different reading frames, which suggests that they could be translated instead of the tORF we were focusing on (tORF_230326, tORF_80553, tORF_102655 and tORF_70680, see Fig. S5 and S8). Eleven of the remaining tORFs display no translation signals and 8 had growth differences in the control conditions so we could not conservatively detect an effect (Fig. S8). Note that among the translated tORFs detected using this approach, 13 were identified only by our custom method for ribosome profiling data.

We next asked if translation was conserved between conditions and strains. We compared the translation of tORFs between the three strains and with the translation pattern observed with ribosome profiling data for the strains in which the DHFR constructs were successful in all three backgrounds. We succeed in transforming five tORFs in all lineages (*SpA*, *SpB* and *SpC*), with translation signals that were consistent with our expression criteria (see Methods). However, we observed that the expression patterns of the tORFs are likely specific to an environment, for instance in SOE medium, tORF_7665 was found to be translated in the *SpC* strain, whereas on the MTX medium, the translation was found only in the *SpB* strain (Fig. S9). Some translation signals were also conserved between strains and conditions, for instance for tORF_14438, which is translated in all three strains in both conditions. These results confirmed the translation detected by ribosomal profiling and indicate that the transcription and translation of tORFs could be highly condition specific, at least for the two conditions considered here (note that the DHFR assay requires a very specific condition). However, the two methods measure slightly different parameters, for instance steady state protein abundance for the DHFR assay and steady state mRNA/ribosome association for the RPF data, which could also contribute to the difference in signals.

## Discussion

To better understand the early stages of *de novo* gene birth, we characterized the properties and turnover of recently evolving iORFs and their putative peptides over short evolutionary time-scales using closely related wild yeast populations. The number of iORFs identified almost doubled when considering within species diversity, which illustrates the possible role of intergenic diversity and the high turnover in providing molecular innovation. Note that we likely underestimate the total number of iORFs segregating in *S. paradoxus* genomes because of our conservative approach to identify a set of unambiguous iORF orthogroups in which we excluded regions too highly divergent that resulted in poor alignments. We focused on ORFs strictly located in intergenic regions but it is important to note that they represent only a subset of non-coding ORFs. Indeed, a recent study has shown that >65% of *de novo* genes arose from transcript isoforms of ancient genes in *Saccharomyces sensus stricto* (Lu et al. 2017). ORFs overlapping known genes (in a different reading frame or in the opposite strand) and pseudogenes may also provide an unneglectable source of ORFs and could be an important contribution to the proteome diversity in wild populations (Ji et al. 2015; Lu et al. 2017; Casola 2018).

The repertoire of iORFs within *S. paradoxus* came from ancient iORFs that are still segregating within *S. paradoxus*, and is regularly supplied with *de novo* iORFs gains. The turnover and retention of iORFs appear at least partly guided by mutation rate variation affecting the number of gains and losses, or by size changes with some larger changes. In addition, longer iORFs were more likely to be submitted to size changes, because of the longer mutational target between the start and stop codons. The iORF turnover rate is lower than the rate of gene duplication or gene loss estimated in yeast (not considering whole genome duplication, (Lynch et al. 2008)) but is high enough to provide closely related lineages with distinct sets of novel ORFs with coding potential.

Among the ~20,000 iORF orthogroups of 60 nt and longer, a small fraction (~2%) showed translation signatures similar to expressed canonical genes in the single condition we tested. Among the 418 tORFS detected using our custom methods, 40% (n=170) were confirmed with the use of another tool (RiboTaper). The detection of translated non canonical ORFs particularly varies depending on the methods, and may lead to only a subset of shared annotated translated ORFs detected, probably due to their generally low translation levels (Xiao et al. 2018). Here, we observed that the use of different methods to detect translation may favor tORFs with different characteristics. For instance, the analysis performed with RiboTaper showed that this tool has more power to detect translation signals on less expressed tORFs, with small initiation peaks. Because we focused on intergenic regions, we gave more importance to translation initiation signals. However, our analysis on expression and sequence properties were robust to translation detection methods.

We observed a stronger post-transcriptional buffering in the tORFs with the highest transcription, reflecting either selection against translation or a lack of selection for optimal translation. This buffering was observed with the use of another ribosome profiling sequencing dataset in *S. cerevisiae* (Fig. S10, McManus et al. 2014). The buffering effect was previously observed in interspecies yeast hybrids, especially for genes that show transcriptional divergence, and was hypothesized to be the result of stabilizing selection on the amount of proteins produced (McManus et al. 2014). In our case, the post-transcription buffering effect is similar between older and younger tORFs, suggesting that selection has instead not been acting or has been too weak to affect this feature.

Consistent with a model in which most tORFs are neutral, the corresponding *de novo* polypeptide properties are on average close to the expectation for random sequences. However, the diversity is large enough that some tORFs have gene-like properties, suggesting a small set of neutrally evolving polypeptides with a potential for new functions. iORF translation signatures (tORFs) were detected for both ancient and recent iORFs and are represented in all conservation groups. This illustrates that there are regular gains and losses of tORFs along the phylogeny. The overall absence of purifying selection acting on tORFs suggests a neutral evolution of most intergenic polypeptides, as observed in rodents (Ruiz-Orera et al. 2018). A study recently found that the expression of random sequences are likely to have an effect on fitness (Neme et al. 2017). By analogy with the fitness effect distribution of new mutations, which are characterized by a large number of mutations of neutral or small effect and few mutations of large effect (Bataillon and Bailey 2014), we hypothesized that only a small fraction of tORFs appearing from random mutations could provide an adaptive advantage strong enough to display a purifying selection signature early after birth. Given this, the resemblance of tORFs to random sequences does not entirely preclude any potential molecular function or fitness effect.

Recently emerging tORFs along terminal branches are more frequent in regions with a higher SNP density, whereas older tORFs tend to be located in slowly evolving regions. This observation suggests variable turnover rates depending on the local mutation rate. Regions with low mutation rates could act as a reservoir of ancient tORFs segregating in the population for a longer time before being lost. On the other hand, mutation hotspots may allow rapid testing of many molecular combinations, which could be advantageous in a changing environment. Most tORFs have a subset of gene-like characteristics, implying that they would require limited refinement by natural selection to acquire new functions. They belong to ancient and recent tORF gain events, suggesting that gene-like characteristics may be conserved over longer evolutionary time scales. These properties could be available immediately for selection to act or when populations are exposed to a changing environment. In addition, even if for a subset of tORFs, the properties are getting closer to gene properties, changes are generally small. This suggests that if they are retained by drift or selection, they provide the raw material to gradually evolve as in the continuum hypothesis (Carvunis et al. 2012). We identified a recently emerging tORF that had several gene-like characteristics, suggesting that it is pre-adapted to be biochemically functional. This example illustrates that the birth of a *de novo* polypeptide may be immediately accompanied with larger gains of gene-like properties, as in the pre-adapted hypothesis (Wilson et al. 2017).

## Material and methods

### Characterization of the intergenic ORFs diversity

We investigated intergenic ORF (iORF) diversity in wild *Saccharomyces paradoxus* populations, which are structured in three main lineages named *SpA*, *SpB* and *SpC* (Charron et al. 2014; Leducq et al. 2016). The wild *S. cerevisiae* strain YPS128 was used in our experiments and the reference S288C (version R64-2-1) was added in our analysis for the functional annotation.

#### Genome assemblies

New genomes assemblies were performed using high-coverage sequencing data from five, ten and nine North American strains belonging to lineages *SpA*, *SpB* and *SpC* respectively 1 (Fig. S1) (Leducq et al. 2016) using IDBA_UD (Peng et al. 2012). For strain YPS128, raw reads were kindly provided by J. Schacherer from the 1002 Yeast Genomes project (Peter et al. 2018). We used the default option for IDBA-UD parameters: a minimum k-mer size of 20 and maximum k-mer size of 100, with 20 increments in each iteration. Scaffolds were then ordered and orientated along a reference genome using ABACAS (Assefa et al. 2009), using the –p nucmer parameter. *S. paradoxus* and *S. cerevisiae* scaffolds were respectively aligned along the reference genome of the CBS432 (Scannell et al. 2011) and S288C (version R64-2-1 from the Saccharomyces Genome Database (https://www.yeastgenome.org/)) strains. Unused scaffolds in the ordering and longer than 200 bp were also conserved in the dataset for further analysis.

#### Identification of homologous intergenic regions

We detected homologous intergenic regions using synteny. Genes were predicted using Augustus (Stanke et al. 2008) with the complete gene model for the species parameter “saccharomyces_cerevisiae_S288C”. Orthologs were annotated using a reciprocal best hit (RBH) approach implemented in SynChro (Drillon et al. 2014) against the reference S288C (version R64-2-1) using a delta parameter of 3. We used RBH gene pairs provided by SynChro and the Clustering methods implemented in Silixx (Miele et al. 2011) to identify conserved orthologs among the 26 genomes. We selected orthologs conserved among all strains and with a conserved order to extract orthologous microsyntenic genomic regions ≥ 100 nt between each pair of genes (Fig. S1).

#### Ancestral reconstructions of intergenic sequences

We reconstructed ancestral genomic sequences of intergenic regions. Because the divergence between strains belonging to the same lineage is low, we chose one strain per lineage to estimate the ancestral intergenic sequences at each divergence node between lineages (Fig. S1 and 1A), that is YPS128 (*S. cerevisiae*), YPS744 (*SpA*), MSH-604 (*SpB*) and MSH-587-1 (*SpC*). The ancestral sequence reconstruction was done using Historian (Holmes 2017), which allows the reconstruction of ancestral indels in addition to nucleotide sequences. Note that indel reconstruction is essential here to not introduce artefactual frameshifts in ancestral iORFs, see below, which depends on the conservation of the same reading frame between the start and the stop codon. Historian was run with a Jukes-Cantor model and using a phylogenetic tree inferred from aligned intergenic sequences by PhyML version 3.0 (Guindon et al. 2010) with the Smart Model Selection (Lefort et al. 2017) and YPS128 as outgroup.

#### iORF annotation and conservation level

Orthologous regions identified between each pair of conserved genes in contemporary strains and their ancestral sequence reconstructions were aligned using Muscle (Edgar 2004) with default parameters. Intergenic regions with a global alignment of less than 50% of identity among strains (including gaps) were removed. We annotated iORFs defined as any sequence between canonical start and stop codons, in the same reading frame and with a minimum size of three codons, using a custom Python script. Because we are working on homologous aligned regions, the presence-absence pattern does not suffer from limitation alignment bias occurring when we are working with short sequences. We extracted a presence/absence matrix based on the exact conservation of the start and the stop codon in the same reading frame (Fig. S1). iORF aligned coordinates were then converted to genomic coordinates on the respective genomes of each strain, and removed if there was any overlap with a known feature annotation, such as rRNA, a tRNA, a ncRNA, a snoRNA, non-conserved genes and pseudogenes annotated on the reference S288C (version R64-2-1 https://www.yeastgenome.org/). Additional masking was performed by removing iORFs i) located in a region with more than 0.6% of sequence identity with *S. cerevisiae* ncRNA or gene (including pseudogenes and excluding dubious ORFs) from the reference genome, or *Saccharomyces kudriavzevii* and *Saccharomyces eubayanus* genes (Zerbino et al. 2018), ii) in a low complexity region identified with repeat masker (http://www.repeatmasker.org/) and iii) when local alignments of iORFs +/-300 bp displayed less than 60% of identity (including gaps). If an iORF overlapped a masked region detected in only one strain, it was removed for all the other strains in order to not introduce presence-absence patterns due to strain specific masking.

iORFs that do not overlap a known feature were then classified according to the conservation level: 1) conserved in both species, 2) specific and conserved within *S. paradoxus*, 3) fixed within lineages and divergent among, 4) specific and fixed in one lineage, 4) polymorphic in at least one lineage (Fig. S1).

For iORFs with a minimum size of 60 nt, we also performed a sequence similarity search against the proteome of NCBI RefSeq database (O'Leary et al. 2016) for 417 species in the reference RefSeq category and the representative fungi RefSeq category (containing 237 fungi species). iORFs with a significant hit (e-value < 10^-3^) were removed to exclude any risks of having an ancient pseudogene. Among the 19,701 iORFs tested, only 12 displayed a significant hit, illustrating the stringency of our thresholds for the iORF annotation and filtering above.

#### Evolutionary history of iORFs

Gain and loss events were inferred by comparing presence/absence patterns between ancestral nodes and actual iORFs. Because the ancestral reconstruction was done using one strain per lineage (see above), polymorphic iORFs absent in all the considered strains have been removed from this analysis. iORFs with no detected ancestral homologs were considered as appearing on terminal branches. We estimated the rate of iORF gain/substitution on each branch as the number of iORF gain divided by the number of substitution (*i.e* branch length × sequence size) and calculated the mean of the four branches. The iORF gain rate per cell per division was estimated by calculating the number of expected substitution per cell per division (from the substitution rate estimated at 0.33x10^-9^ per site per cell division by Lynch et al. (2008), multiplied by the iORF gain rate per substitution.

The evolution of iORF sizes was inferred by connecting iORFs with their ancestral homologs along the phylogeny if they shared the same start and/or stop position on aligned intergenic sequences. iORF sizes of two connected iORFs may be conserved if there are no changes, an increase or a decrease if there are connected only by the same start or stop position because the position of the other extremity of the iORFs changed.

### Ribosome profiling and mRNA sequencing libraries

Ribosome profiling and mRNA sequencing experiments were conducted with the strains YPS128 (*S. cerevisiae)* (Sniegowski et al. 2002) and YPS744 (*S. paradoxus*), MSH604 (*S. paradoxus*) and MSH587 (*S. paradoxus*) belonging respectively to groups *SpA*, *SpB* and *SpC* according to Leducq et al. (2016). We prepared two replicates per strain and library type. The protocol is described in supplementary methods. Briefly, strains were grown in SOE (Synthetic Oak Exudate) medium (Murphy et al. 2006). Ribosome profiling footprints were purified using the protocol described in Baudin-Baillieu et al. (2016) with modifications (see supplementary methods). The rRNA was depleted in purified ribosome footprints and total mRNA samples using the Ribo-Zero Gold rRNA Removal Kit for yeast (Illumina) according to the manufacturer’s instructions. Ribosome profiling and total mRNA libraries were constructed using the TruSeq Ribo Profile kit for yeast (illumina), using manufacturer’s instructions starting from fragmentation and end repair step. Libraries were sequenced with Illumina HiSeq 2500 at The Genome Quebec Innovation Center (Montreal, Canada).

### Detection of translated iORFs

Both total RNA and ribosome profiling sequencing libraries were processed using the same procedure. Raw sequences were trimmed of 3’ adapters using CUTADAPT (Martin 2011). For RPF data, reads with lengths of 27–33 nucleotides were retained for further analysis as this size is most likely to represent footprinted fragments. For mRNA, reads with lengths of 27–40 nucleotides were retained. Adapter trimmed reads were aligned to the respective genome of each sample using Bowtie version 1.1.2 (Langmead et al. 2009) with parameters –best – chunkmbs 500.

We used ribosome profiling reads to identify translated iORFs using a custom method. This analysis was performed on iORFs longer or equal to 60 nucleotides to detect translation signatures and codon periodicity on at least 20 codons. Annotated iORFs may be overlapping because of the three possible reading frames for each strand. Ribosomal speed differences during translation cause an accumulation of ribosome footprints at specific positions within a gene (Ingolia 2016). We used ribosome profiling read density, which is typically characterized by a strong initiation peak located at the start codon followed by a codon periodicity at each codon, to detect the translated iORF among overlapping ones. For each strain, we performed a metagene analysis at the start codon region of iORFs and annotated conserved genes to detect the p-site offset for each read length between 28 and 33 nt. Because the ribosome profiling density pattern is stronger in highly translated regions, metagene analyses were done using the two replicates of each strain pooled in one coverage file. Ribosome footprints were mapped to their 5’ ends, and the distance between the largest peak upstream of the start codon and the start codon itself is taken to be the P-site offset per read length. When comparing annotated genes and iORFs, we obtained similar P-site offset estimates per read length, which were used for next analysis. We then extracted the aligned read densities, subtracted by the P-offset estimates, per iORF or gene for next analyses. Metagene analyses were performed using the metagene, psite and get_count_vectors scripts from the Plastid package (Dunn and Weissman 2016), metagene figures were done using R scripts (R Core Team 2013).

We identified translation initiation signals from ribosome profiling per base read densities, by detecting peaks at the start codon using a custom R script. We defined three precision levels of peak initiation: ‘p3’ if the highest peak is located at the first nucleotide of the start codon, ‘p2’ there is a peak at the first position of the start codon and ‘p1’ if there is a peak at the first position of the start codon +/-1 nucleotide. A minimum of five reads was required for peak detection. Read phasing was estimated by counting the number of aligned reads at the first, second or the third position for all codons, excluding the first one, of the considered iORF or gene, to test for a significant deviation from expected ratio with no periodicity, that is 1/3 of each, with a binomial test. We applied an FDR correction for multiple testing. A minimum of 15 reads was required for phasing detection.

iORF families or genes with an initiation peak and a significant periodicity, *i.e.* a FDR corrected p-value < 0.05, in at least one strain were considered as translated and named tORFs.

We detected translation signature using the RiboTaper software (Calviello et al. 2016). We used read lengths for which we obtained the best in frame phasing with annotated genes according to quality check plots provided by RiboTaper, and which are 30-31 nt for *SpA*, 30-32 for *SpB* and 31-32 for *SpC*, and a P-offset of 13.

### Differential expression analysis

Reads were strand-specifically mapped to tORFs and conserved genes using the coverageBed command from the bedTools package version 2.26.0 (Quinlan and Hall 2010), with parameter −s (Supplemental Table S3). We then examined significant tORF expression changes between strains. The differential expression analysis was performed using DESeq2 (Love et al. 2014). Significant differences were identified using 5% FDR and 2-fold magnitude. We identified lineage specific expression increase when the expression of the tORFs in the considered lineage was significantly more expressed than the others strains in all pairwise comparisons. For *SpB-SpC* increase, we selected tORFs when *SpB* and *SpC* strains were both more expressed than YPS128 and *SpA,* and *S. paradoxus* increase when all *S. paradoxus* lineages were more expressed than YPS128.

For the visualization of tORF coverages (Fig. 5 and Fig. S6), we extracted the per base coverage on the same strand using the genomecov command from the bedTools package version 2.26.0 (Quinlan and Hall 2010). The normalization was performed by dividing the perbase coverage of each library with the size factors estimated with DESeq2 (Love et al. 2014).

### Strain construction for in vivo translation confirmation

45 tORFs along with 12 canonical genes (Supplemental Table S5) were tagged with a modified full-length DHFR — a marker that gives resistance to methotrexate (Tarassov et al. 2008) — in frame and out of frame (as a control). The tORFs were chosen due to their strong translation signature differences between lineages as found by the differential expression analysis with ribosome profiling. If the tORF is indeed expressed, in-frame DHFR-tagged strains should grow in medium supplemented with methotrexate. This complements the ribosome profiling as an *in vivo* confirmation of tORF expression.

DHFR along with a HPH resistance module (on a pAG32-DHFR1,2-3 (synthesized by Synbio Tech, New Jersey, USA)) were PCR amplified (Kapa Hifi DNA polymerase – Kapa Biosystems Inc., Wilmington, USA) using primers that, at each end, added homology regions flanking the stop codon of the tORF of interest (Supplemental Table S4). Forward primers were flush with the stop codon for the in frame integration, and −2bp for the out of frame one (figure 6A). To fuse the DHFR with the tORFs, 8 µl of the PCR products were then used for transformations in *SpA* (YPS744), *SpB* (MSH604) and *SpC* (MSH587-1) (only *SpC* for the canonical genes) according to the method described in (Bleuven et al. 2018).

Successful transformations were confirmed by growth on YPD + 250 µg/ml hygromycin B (HYG) + 100 µg/ml Nourseothricin (NAT) and by PCR amplification of the region containing the tORF tagged with DHFR.

### Phenotyping of DHFR-tagged strains

Transformed strains were incubated at 30°C in 2ml 96-deepwell plates containing 1ml of liquid YPD+HYG+NAT for 24h. From there, different 96-arrays were made and the strains were printed onto solid YPD+HYG+NAT plates (omnitrays) using a robotic platform (BM5-SC1, S&P Robotics Inc.) with appropriate pin tools (96, 384 and 1536). Plates were incubated two days at 30°C. The solid media 96-arrays were pinned into 384-arrays and then, into the 1536-array with which the phenotyping was done. The final 1536-plate was then replicated into the same format on a second YPD+HYG+NAT plate to get more uniformly sized colonies. Plates were incubated two days at 30°C between each steps. All strains were present in five or six replicates. To avoid positional effects of the plate borders, the two outer rows and columns were filled with a control strain (BY4743 LSM8-DHFR[1,2]/CDC39-DHFR[3]).

To test for methotrexate resistance, all strains were then transferred to DMSO (control) and MTX DHFR PCA media (0.67% yeast nitrogen base without amino acids and without ammonium sulfate, 2% glucose, 2.5% noble agar, drop-out without adenine, methionine and lysine, and 200 µg/mL methotrexate (MTX) diluted in DMSO (or only DMSO in the control medium)). Plates were incubated at 30°C for four days, after which a second round of MTX selection was performed. Plates were incubated at 30°C for another four days. Images were taken with an EOS Rebel T5i camera (Canon) every two hours during the entire course of the experiment. Incubation and imaging was performed in a spImager custom platform (S&P Robotics Inc.).

Images were processed using the gitter.batch function in the R package Gitter (Wagih, Parts 2014 – Version 1.1.1). The last image of each experiment was used as a reference image to ensure accurate identification of colonies at early timepoints. The size after 60 hours of growth (the 30th image) was extracted and the median was calculated for the replicates, these values are the base for figure 6B (Supplemental Table S6). In-frame and out of frame strains were phenotyped together on the same plate to alleviate batch effects. Translation was detected i) when we observed colony size differences between in-frame and out of frame constructions on MTX medium with a student t-test (p-value < 0.05), and ii) if both positive controls display colony sizes of more than 1000 and with similar growth for both controls.

Some of the observed results were confirmed by measuring cell growth in a spot-dilution assay. Briefly, precultures of cells expressing DHFR fused to tORFs of interest were adjusted to an OD600/mL of 1 in water. 5-fold serial dilutions were performed and 6 µL of each dilution were spotted on DMSO and MTX DHFR PCA media. Plates were incubated for five days at 30°C and imaged each day with an EOS Rebel T3i camera (Canon).

### Expression and sequence properties

Normalized read counts for ribosome profiling and total mRNA samples were extracted with DESeq2 software (Love et al. 2014) and we calculated the mean of the two replicates per library type. Translation efficiency (TE) was calculated as the ratio of RPF over total mRNA normalized read counts on the first 60 nt. We excluded tORFs and genes with less than 10 total RNA reads in the first 60 nt for the TE calculation. Slope differences between genes and tORFs were tested with an ANCOVA. We confirmed the buffering effect on tORFs annotated in the *S. cerevisiae* reference strain S288C with ribosome profiling and RNA sequencing data obtained in (McManus et al. 2014) (Fig. S10).

The intrinsic disorder was calculated for genes and intergenic tORFs using IUPRED (Dosztanyi et al. 2005). The SNP rate was calculated for each syntenic intergenic region by dividing the total number of intergenic SNPs in *S. paradoxus* alignments, by the total number of nucleotides in the region, as in Agier and Fischer (2012) study for intergenic sequences. We used the *codeml* program from the PAML package version 4.7 (Yang 2007) to estimate the likelihood of the dN/dS ratio, using the same procedure as employed by Carvunis et al. (2012) with codon model 0.

All analyses were conducted and figures were created using python and R (R Core Team 2013).

## Data access

High-throughput sequencing data generated in this study have been submitted to the NCBI BioProject database (https://www.ncbi.nlm.nih.gov/bioproject) under accession number PRJNA400476. Assemblies and annotations are available at https://landrylab.ibis.ulaval.ca/?page_id=2211.

## Acknowledgments

We thank G. Charron and the IBIS sequencing platform (B. Boyle) for technical help and A.R Carvunis, R. Dandage and the reviewers for comments on the manuscript. This project was funded by a FRQNT Team grant to C.R.L and Xavier Roucou and NSERC discovery grant to C.R.L. C.R.L. holds the Canada Research Chair in Evolutionary Cell and Systems Biology.

## Author contributions

E.D and C.R.L conceived the project. E.D, O.N, I.H, and I.G.A designed ribosome profiling experiments. E.D, I.G.A and I.H performed ribosome profiling experiments. A.K.D, J.H, I.G.A and C.R.L designed and performed functional validation experiments. E.D performed the bioinformatics analyses with helpful advices from L.N.T, C.R.L and O.N. E.D wrote the manuscript with revisions from all authors.

