## Supplementary information for "The high turnover of ribosome-associated transcripts from *de novo* ORFs produces gene-like characteristics available for *de novo* gene emergence in wild yeast populations"

- Supplementary analysis (page 2)
- Supplementary methods (page 6)
- Supplementary figures (page 10)
- Supplementary tables (page 32)
- References (page 32)

**Supplementary analysis**

A large number of intergenic ORFs segregate in wild *S. paradoxus* populations

We characterized iORF diversity in wild *S. paradoxus* populations. We used genomes from 24 strains that are structured in three main lineages named *SpA*, *SpB* and *SpC* (Charron et al. 2014; Leducq et al. 2016). Two *S. cerevisiae* strains were included as outgroups: the wild isolate YPS128 (Sniegowski et al. 2002; Peter et al. 2018) and the reference strain S288C. These lineages cover different levels of nucleotide divergence, ranging from ~ 13 % between *S. cerevisiae* and *S. paradoxus* to ~2.27 % between the two closest *SpB* and *SpC* lineages (Kellis et al. 2003; Leducq et al. 2016). We used microsynteny to identify and align orthologous non-genic regions between pairs of conserved annotated genes (Fig. S1 and Methods). We identified 3,781 orthologous sets of intergenic sequences representing a total of ~ 2 Mb, with a median size of 381 bp (Fig. S1 and S12). iORFs were annotated on aligned sequences using a method similar to the one employed by Carvunis et al. (2012), that is the first start and stop codons in the same reading frame not overlapping with known features, regardless of the strand, and with no minimum size. We then classified iORFs according to their conservation level among strains (Fig. S1 and Methods). Because the annotation was performed on aligned sequences, we could precisely detect the presence/absence of orthologous iORFs among strains, based on the conservation of an iORF with the same start and stop positions without disruptive mutations in between. We used *S. cerevisiae* as an outgroup and removed iORFs present only in this species in order to focus on *S. paradoxus* diversity. However, we conserved iORFs present both in *S. cerevisiae* and in at least one *S. paradoxus* strain to consider the inter-species conservation.

We annotated 34,216 to 34,503 iORFs per *S. paradoxus* strain, for a total of 64,225 orthogroups annotated at least in one *S. paradoxus* strain (Table 1). This represents a density of about 17 iORFs per Kb. The iORF set shows about 6 % conserved among *S. cerevisiae* and *S. paradoxus* strains, and 15 % specific and fixed within *S. paradoxus.* The remaining 79 % are still segregating within *S. paradoxus* (Fig. S13A, S13B and Table 1).

To understand how iORF diversity changes over a short evolutionary time scale, we estimated the age of iORFs and their turnover using ancestral sequence reconstruction (Fig. S13C) (see Methods). Because polymorphism within lineages (*SpA*, *SpB* or *SpC*) (Leducq et al. 2016) may affect the topology of the phylogeny (although most diversity in this group is among lineages), we used only one strain per lineage (YPS128 (*S. cerevisiae*), YPS744 (*SpA*), MSH-604 (*SpB*) and MSH-587-1 (*SpC*)) to reconstruct ancestral sequences at two divergence nodes that we labeled N1 for *SpB-SpC* divergence and N2 for *SpA-SpB/C* divergence. These strains contain 58,952 iORF orthogroups after removing the polymorphic iORFs that are absent in all the four selected strains. Reconstructed sequences were included in intergenic alignments of actual strains and were used to detect the presence or absence ancestral iORFs at each node (Fig. S1, S13C and Methods).

We estimated the age of the 58,952 iORFs and annotated the 2,291 iORFs detected only in ancestral sequences. 55 % of iORFs were present at N2 (the oldest age category) and are represented in each conservation group depending on iORF loss events occurring after N2 (Fig. S13D and Table 2). We observed a continuous emergence of iORFs with 6,782 gains between N2 and N1, and 5,324 to 8,454 along terminal branches. As expected, the number of iORF gains or losses is correlated and increases with branch length (Fig. S13E). We estimated a rate of emergence and loss at respectively 0.28 +/- 0.01 and 0.27 +/-0.008 ORFs per nucleotide substitution. An ORF is on average gained or lost at every 3.5 substitutions. The *de novo* ORF gain rate, estimated at around $1.1\times10^{-3}$ ORFs per genome per cell division, is about one order of magnitude smaller than the rate of gene duplication in *S. cerevisiae* estimated at $1.9 \times 10^{-2}$ genes per genome per cell division (Lynch et al. 2008).

We considered that iORFs with no detected ancestral homologs appeared on terminal branches. Among them, 91 to 93 % are present only in one lineage, which is consistent with the expected conservation pattern for recently emerging iORFs (Table 2). The absence of ancestral homologs for the remaining 7 to 9% iORFs present in more than one lineage can be due to convergence on terminal branches, made possible by the relatively high turnover rate. Convergence events may particularly occur if two lineages acquire independently small indels, not necessarily at the same position but in the same iORF, leading to the same frameshift and resulting in stop codon changes. To estimate the expected frequency of such convergence events, we counted the number of iORFs for which the observed SNPs or indels lead to convergence in the alignment of 133 iORFs present in both *SpB* and *SpC* lineages, with no detected ancestors. We observed that ~30 % of patterns of convergence were indeed due of convergent mutations. Finally, regions with a higher rate of evolution may more likely lead to ancestral sequence reconstruction errors and to a small overestimation of the gain rate but this effect should be negligible because of the small number of iORFs with ambiguous age estimation.

As previously observed, iORFs tend to be small with a median value of 43 bp compared to known genes in the reference *S. cerevisiae* (median gene size of 1,287 bp) (Fig. S14A). Each conservation group also contains iORFs longer than the smallest annotated genes in *S. cerevisiae*, revealing an extended set of iORFs with coding potential. In our study, overlapping iORFs between strains, sharing the same start and a different stop position (or the reciprocal) were classified as different orthogroups because of their different sizes. We investigated the evolution of iORF sizes along the phylogeny, by connecting overlapping iORF orthogroups in actual strains with their ancestral homologs based on the conservation of their start and/or stop positions (Fig. S14B). The majority of iORF orthogroups (65%) were conserved until N2 (Fig. S14B). We identified 19% of iORFs successively connected to N1 and N2 by one or two size changes along the phylogeny (Fig. S14B). Note that a size change is considered as a loss event generally accompanied by the gain of another iORF, which is consistent with the similar gain and loss rates estimated. iORFs detected only on terminal branches with no ‘connected’ ancestral homologs tend to display intermediate iORF size values compared to iORFs of conserved size and iORFs resulting from size changes (Fig. S14C).

Size changes are also mainly small even if some extreme cases are observed (Fig. S14D-E). Compared to smaller iORFs, longer iORFs are less conserved and more submitted to size changes (Chi-square test, p-value $< 2.2 \times 10^{-16}$, Fig. S14C and S14F), which might be explained by the higher turnover rate of longer sequences, which are a larger target for mutation accumulation. Longer iORFs also tend to decrease, which might be due to a higher probability of acquiring a disruptive mutation resulting in a size decrease, and intergenic size constrains limiting the maximum iORF sizes (Fig. S12 and S14E).

### Supplementary methods

Ribosome profiling and mRNA sequencing libraries

*Polysome extract preparation*

Ribosome profiling and mRNA sequencing experiments were conducted with the strains YPS128 (*S. cerevisiae)* (Sniegowski et al. 2002), YPS744 (*S. paradoxus*), MSH604 (*S. paradoxus*) and MSH587 (*S. paradoxus*) belonging respectively to groups *SpA*, *SpB* and *SpC* according to Leducq et al. (2016). All strains were diploid. For *S. paradoxus*, we constructed homozygous diploids from haploid heterothallic strains containing a resistance cassette (Nourseothricine or Hygromycine B depending of the mating type) at the *HO* locus. Constructions and crosses were performed according to the protocol described in Leducq et al. (2016). Resulting diploid cells containing the two resistance cassettes were selected on solid YPD (Yeast Peptone Dextrose) medium containing100 ug/ml of Nourseothricine and 250 ug/ml of Hygromycine B.

Strains were grown overnight in 50 mL of SOE (Synthetic Oak Exudate) medium (Murphy et al. 2006), at 30°C with shaking at 250 rpm. These pre-cultures were used to inoculate a 30°C pre-warm 750 mL SOE medium at an initial $OD_{600}$ of ~0.03, and grown to an $OD_{600}$ between 0.6 to 0.7, at 30°C and shaking at 250 rpm. We choose the SOE medium to be closed to natural conditions in which *de novo* genes could emerge in wild yeast strains. Cultures were treated with cycloheximide (50 ug/mL final) for 5 minutes and cells were rapidly collected by vacuum filtration using a 90 mm cellulose nitrate filter with a 0.45 mm pore size and a fritted glass support.  Cells were resuspended on ice in 2.5 mL of polysome lysis Buffer (10 mM Tris-HCL pH 7.4, 100 mM NaCl, 30 mM MgCl2, 50 ug/mL cycloheximide).  The slurry was then pipetted and frozen by fractions of ~ 20 ul in liquid nitrogen, and stored at -80°C. The resulting cryogenized mix was grinded in a MixerMill 400 (RETSCH) for 15 cycles of 2mn at 30 Hz, with chilling in liquid nitrogen between each cycle. The powder was gently thawed in the open grinding chamber at room temperature to collect the lysate which was cleared by two rounds of centrifugation for 5 minutes at

3000xg at 4°C, followed by one round of high speed centrifugation for 10 min at 20,000xg at 4°C. The middle layer was quantified by $OD_{260}$ measurement via Nanodrop and samples above 200 $OD_{260}$/mL were diluted to ~200 $OD_{260}$/mL in lysis buffer. Cell lysates were divided into 250 ul aliquots of 30 to 50 $OD_{260}$ each, that were flash frozen in liquid nitrogen and stored at -80°C. For each lysate, one aliquot of 250 ul was conserved for direct total mRNA extraction, the other aliquots were pooled for ribosome footprint isolation.

*Isolation and purification of ribosome footprints*

Cell lysate, corresponding to 50 to 100 $OD_{260}$, were digested with 15 U of RNAse I (Ambion) per $OD_{260}$ for 60 minutes at 25 °C with shaking. The digestion was stopped by adding 200 U of Superase-in (AMBION). The digested products were loaded on a 24% sucrose cushion (50 mM Tris-acetate pH 7.6, 50 mM $NH_4Cl$, 12 mM $MgCl_2$, 1 mM DTT) and centrifuged at 4°C 100,000 rpm in a TLa110 rotor for 2h15. Pellets were washed two times with lysis buffer and resuspended in 500 uL of polysome lysis buffer. The extract was treated with DNAse I using the manufacturer's instructions (Truseq Ribo profile illumina kit for yeast). RNA was then extracted using acid-phenol-chloroform extraction protocol, and precipitate overnight at -20 °C with 0.1 Volume of sodium acetate 3M, pH 5.2 and 3 volumes of EtOH 100%. Samples were centrifuged at 4°C for 20 minutes at 10,000 g and pellets were resuspended in 75 ul of RNAse free $H_20$ supplied with 1 U/ul of Superase-in (AMBION). RNA concentration was measured at 260 nm, resulting in a final amount of ~300 to 1000 ng of digested RNA. RNA fragments were separated by electrophoresis on a denaturing 17% PAGE gel with heating at 60 °C at 200V for ~8 hours. A mix of 28 and 34 nt RNA markers, oNTI199 and oNTI34ARN (Ingolia et al. 2012), was loaded at both extremities of the gel. The gel was stained with SYBR Gold according to the manufacturer's instructions and the region corresponding to the 28 marker was excised. Gel slices were disrupted through needle holes in a 0.5 mL centrifuge tubes nested in a 1.5 mL tube by maximum speed centrifugation. RNA was eluted overnight at 4°C with gentle rotation in an elution buffer (300 mM NaOAc pH 5.5, 1 mM EDTA). The slurry was loaded on SpinX cellulose acetate filter to recover the eluted RNA cleared of gel fragments. The RNA was precipitate overnight at -20°C in ethanol with 0.3 M sodium acetate and 20 ug of glycogen. Samples were centrifuged at 4°C for 30 minutes at maximum speed and pellets were resuspended in 25 ul of nuclease-free water supplemented with 0.1 U/mL of Superase-in (AMBION). These samples contain purified ribosome footprints. Total mRNA was extracted using the same acid-phenol-chloroform extraction protocol as for ribosome footprints samples. Purified ribosome footprints and total RNA were then quantified by fluorescence (Quant-it RNA assay kit, thermofisher) and stored at -20°C.

*Library preparation*

The rRNA was depleted in purified ribosome footprints and total mRNA samples using the Ribo-Zero Gold rRNA Removal Kit for yeast (Illumina) according to the manufacturer’s instructions. Ribo-Zero treated RNAs were then purified by overnight ethanol precipitation. Ribosome profiling and total mRNA libraries were constructed using the TruSeq Ribo Profile kit for yeast (illumina), using manufacturer’s instructions starting from Fragmentation and end repair step. Circularized cDNA templates were amplified by 11 cycles of PCR using Phusion-polymerase (New England Biolabs), with primers incorporating barcoded Illumina TruSeq library sequences, according to TruSeq Ribo Profile kit for yeast (illumina). The resulting PCR products were loaded onto a 8% native polyacrylamide gel in TBE and purified using the PCR purification protocol provided in the TruSeq Ribo Profile kit for yeast (illumine). The quality and size of the purified PCR products were assessed using an Agilent HS bioanalyzer. Libraries were quantified by fluorescence using the Quant-iT PicoGreen dsDNA Assay Kit (ThermoFisher). The 8 total RNA libraries were pooled in one bulk for sequencing. RPF libraries were pooled in 2 bulks, one for the first replicate of the 4 strains, and a bulk for the second replicate of the 4 strains. Libraries were sequenced on an Illumina HiSeq 2500 at The Genome Quebec Innovation

Center. The total RNA bulk was loaded onto 5 lanes and RPF bulks were loaded onto 4 lanes each.

### Supplementary figures

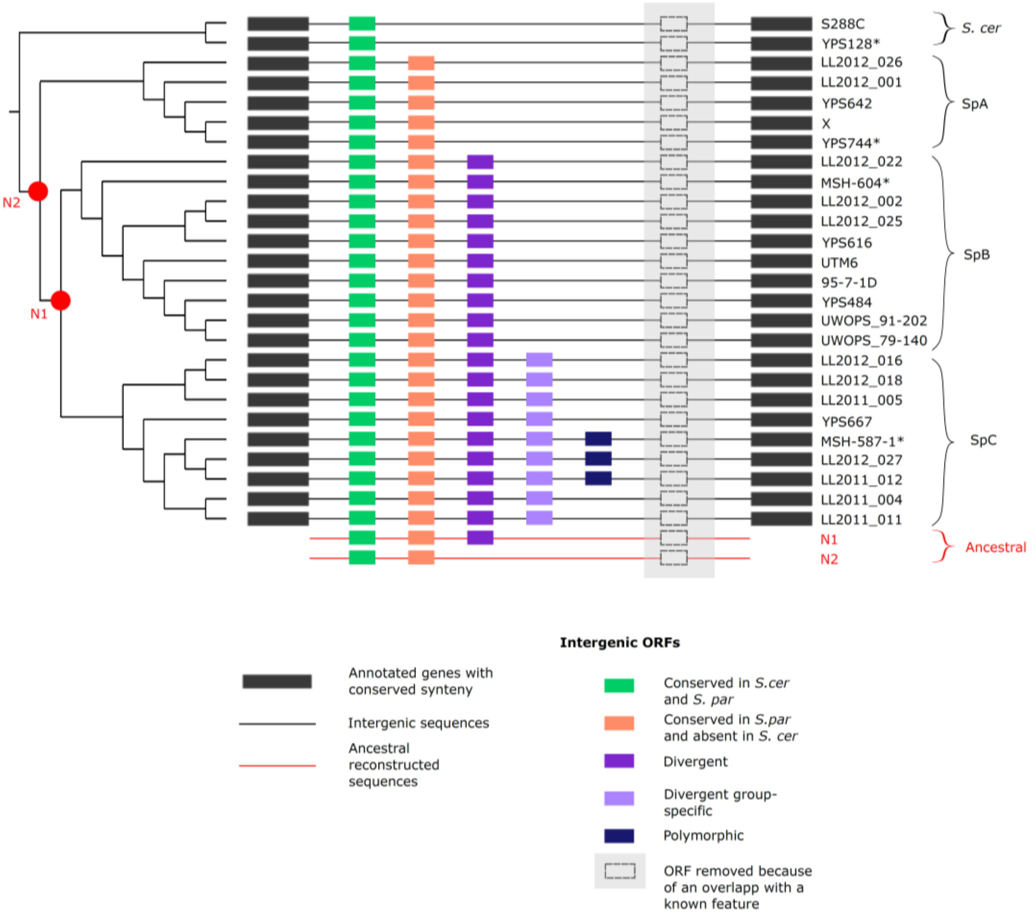


**Figure S1. Identification and annotation of iORFs.** Genes with conserved synteny were used as anchor to align non-genic sequences for each pair. iORFs were annotated on intergenic aligned sequences and clustered as orthogroups based on the conservation of the positions of their start and stop codons aligned with no disruptive mutation within. iORFs displaying a sequence similarity or an overlap with a known feature were removed from all strains. Strains used for ribosome profiling experiments are marked with *.

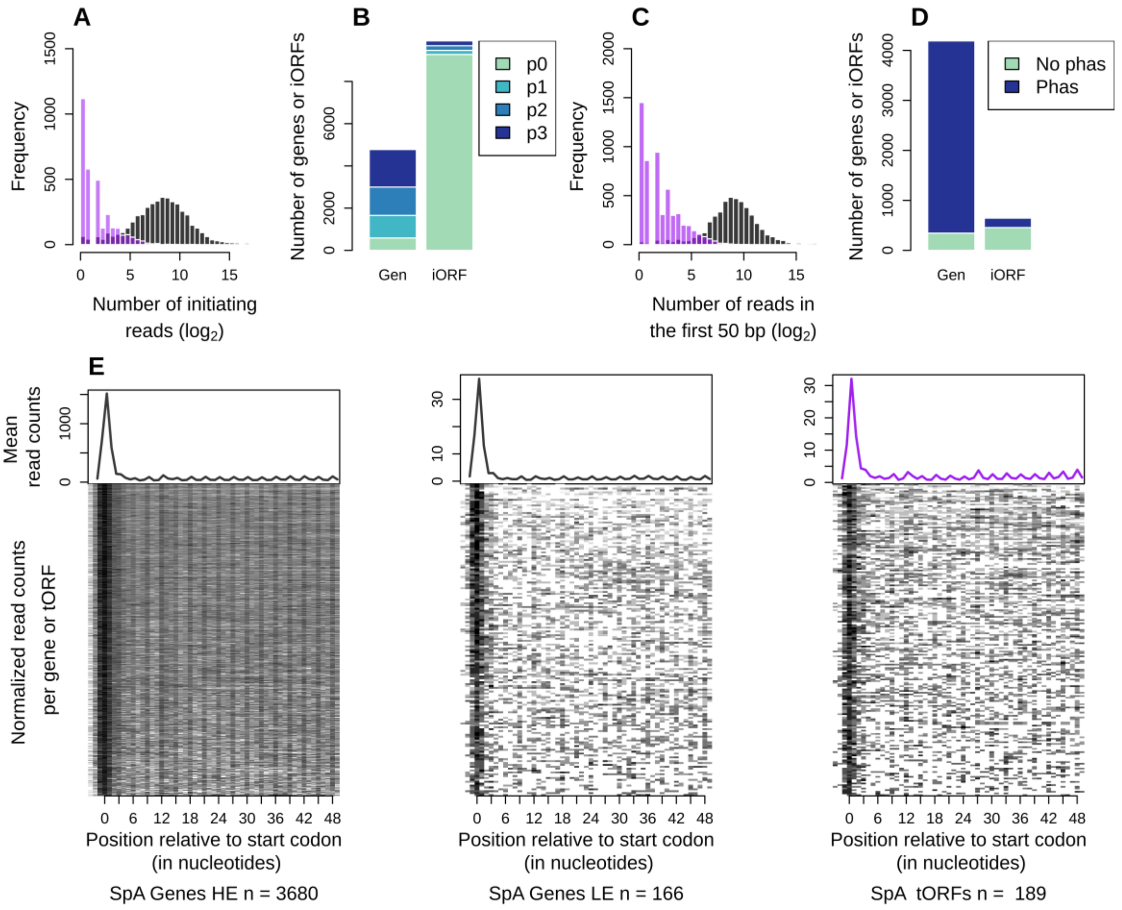


**Figure S2. Detection of iORF translation signatures.** Results for *SpA* and *SpB* strains (see Fig. 2 for *SpC* results). **A)** Distribution of ribosome profiling read counts for genes (in grey) and iORFs (in purple) at the start codon position. **B)** Proportions of genes (Gen) or iORFs with a detected initiating peak at the start codon position. Peaks are colored according to the precision of the detection (see Methods), from the most precise (p3) to the less precise (p1). No peak detection is in green (p0). **C)** Distribution of ribosome profiling read counts in the first 50 nt of iORFs excluding the start codon. **D)** Proportions of genes or iORFs with a significant codon periodicity (in blue) among genes and iORFs with a detected initiation peak. No peak detection is in green. **E)** Metagene for significantly translated and highly (HE) or lowly (LE) expressed genes in grey, and intergenic tORFs in purple. The mean of 5' read counts is plotted along the position relative to the start codon for significantly translated genes or tORFs. Lines of the matrix indicate the normalize coverage of all genes or tORFs with significant signature of translation.

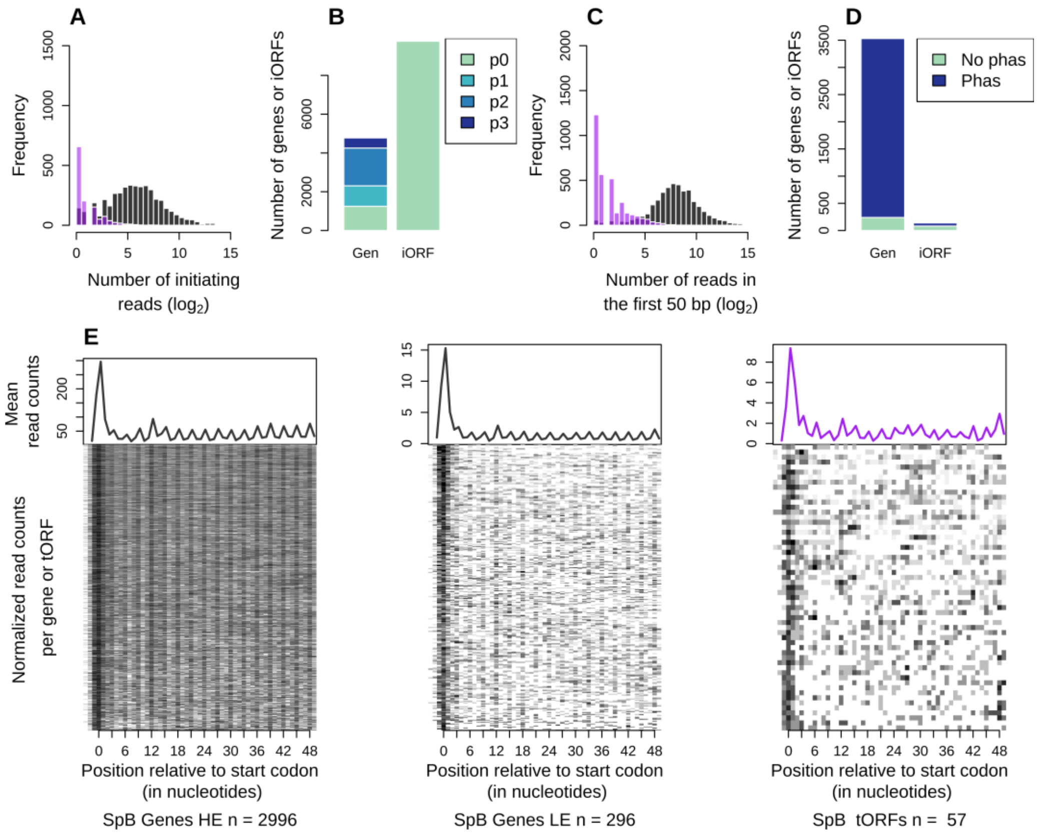


**Figure S2. Continued.**

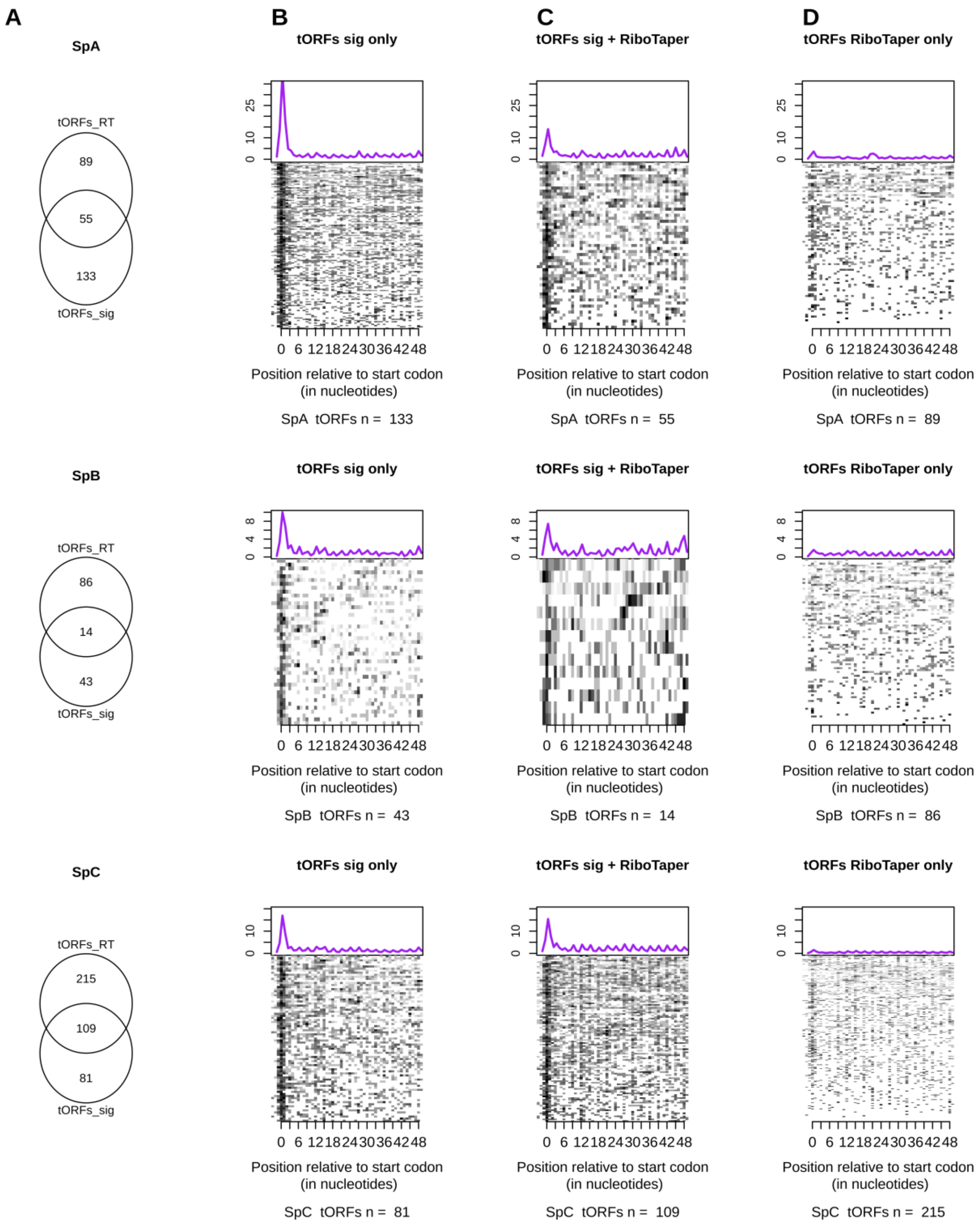


**Figure S3. Translation profiles of tORFs with two detection methods.** tORFs_sig and tORFs_RT respectively represents tORFs detected with our custom method and RiboTaper. **A)** Venn diagram showing the number of detected tORFs depending on the method per haplotype. **B-D)** Metagene analysis for tORFs detected **B)** only with our method, **C)** with both methods and **D)** with RiboTaper only. The mean of 5' read counts is plotted along the position relative to the start codon for significantly translated genes or tORFs. The lines of the matrix indicate the normalize coverage of genes or tORFs with significant signature of translation, with one feature per line.

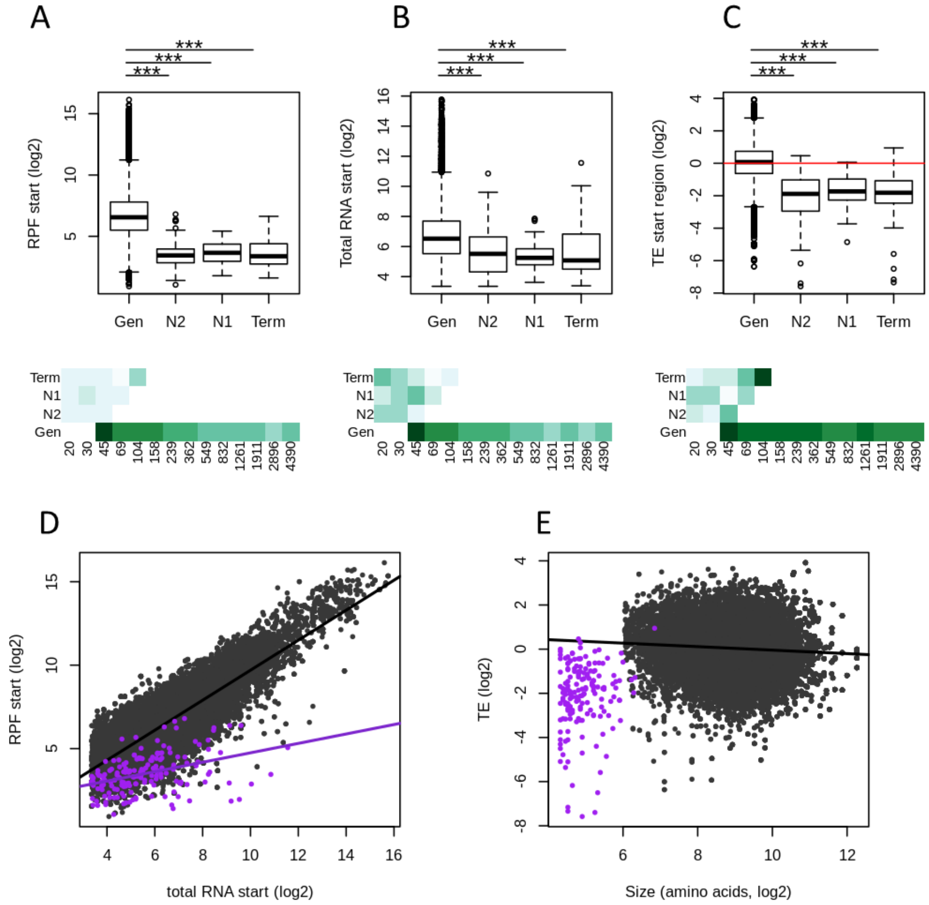


**Figure S4. Expression properties of tORFs detected with RiboTaper (see Fig. 3 for tORFs detected with our custom method). A-C)** Ribosome profiling (RPF), total RNA and translation efficiency (TE) - read counts in the first 60 nt, normalized to correct for library size differences in log2 - are displayed for genes (Gen) and tORFs according on their age (N2, N1 and Term). Significant differences in pairwise comparisons are displayed above each plot (Wilcoxon test, *** for p-values < 0.001, ** for p-values < 0.01 and * for p-values < 0.05). Mean estimates per size range are colored by green intensities (from pale for low values to dark green for high values) below. **D)** RPF plotted as a function of total RNA for tORFs in purple, or genes in grey**. E)** TE plotted as a function of tORF or gene sizes (number of amino acid residues in $log_2$). Regression lines are plotted for significant Spearman correlations (p-values < 0.05). Expression levels were calculated using the mean of the two replicates.

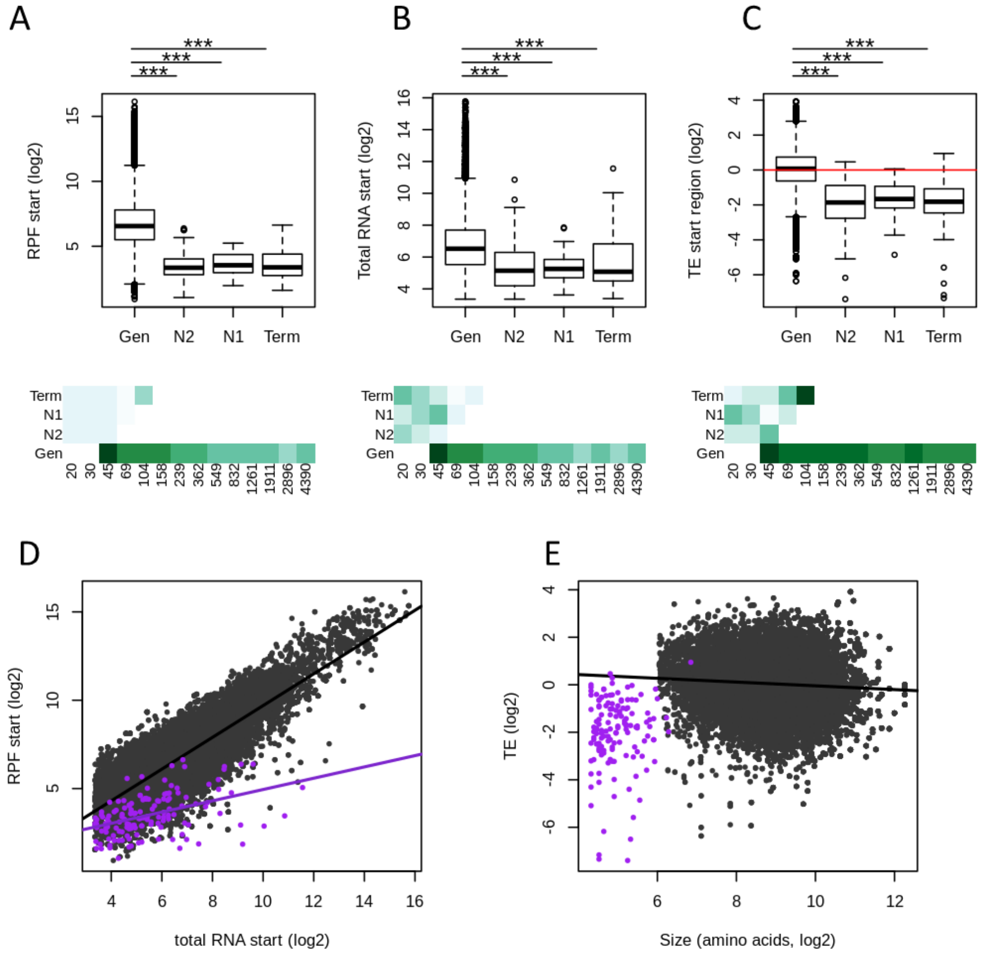


**Figure S4. Continued. Expression properties of tORFs detected with both our custom analyses and RiboTaper.**

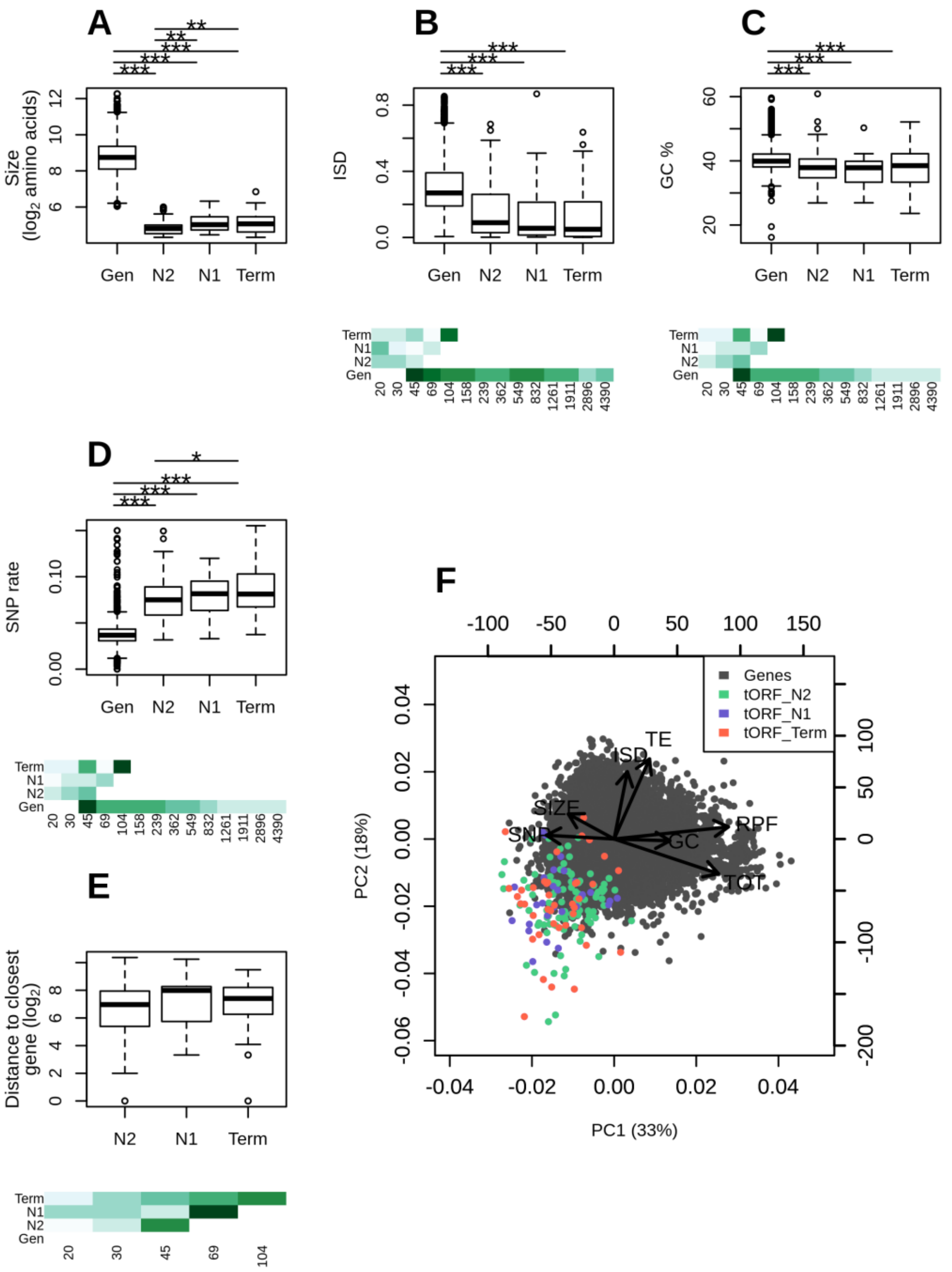


**Figure S5. Sequence properties of tORFs detected with RiboTaper (see Fig. 4 for tORFs detected with our custom method). A-E)** Sizes ($log_2$ number of residues), mean disorder (ISD), GC content (%), SNP density and distance to the closest gene are displayed for genes and tORFs as a function of age (N2, N1 and Term). Pairwise significant differences are displayed above each plot (Wilcoxon test, *** for p-values < 0.001, ** for p-values < 0.01 and * for p-values <0.05). Mean estimates per size ranges are colored with green intensities (from pale for low values to dark green for high values) below. **F)** Principal component analysis using the number of residues (SIZE in $log_2$), ribosome profiling (RPF), total RNA (TOT) and translation efficiency (TE) (as read counts in the first 60 nt normalized to correct for library size differences and in $log_2$), intrinsic disorder (ISD), the GC content and SNP density (SNP). tORFs are colored according to their age.

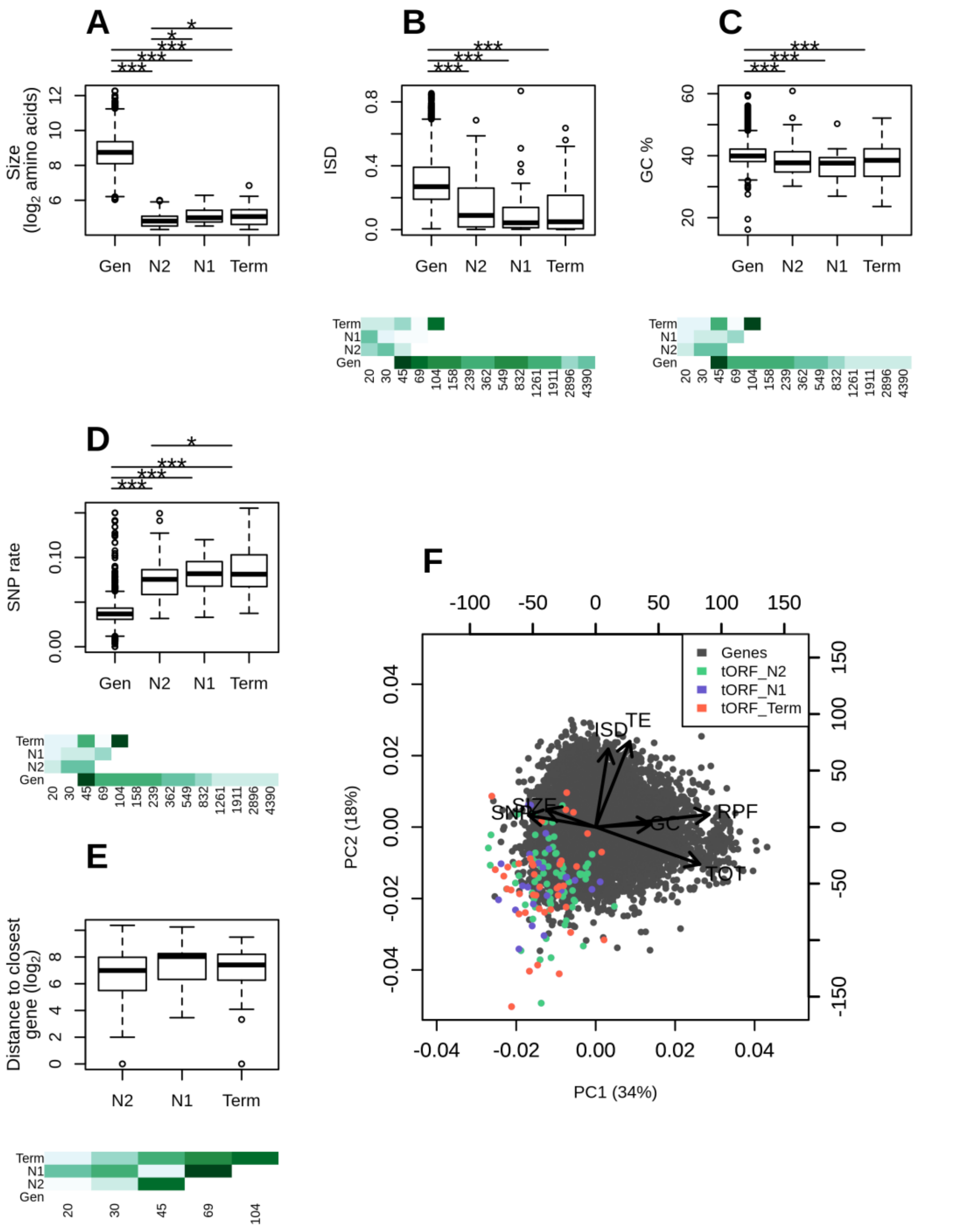


**Figure S5. Continued. Sequence properties of tORFs detected with both our custom analyses and RiboTaper.**

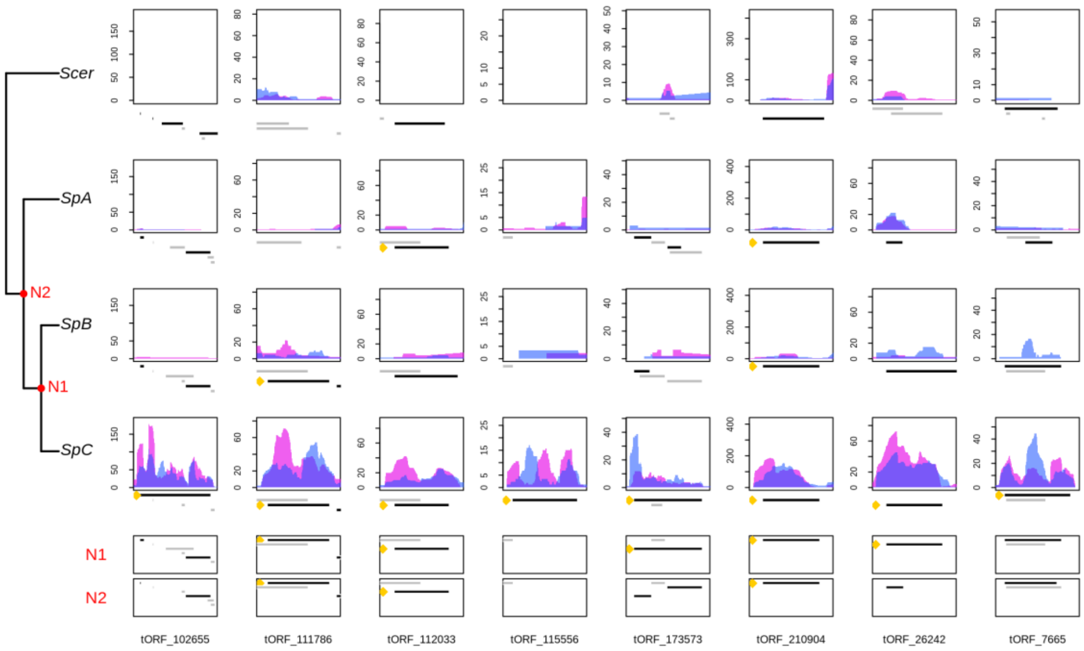


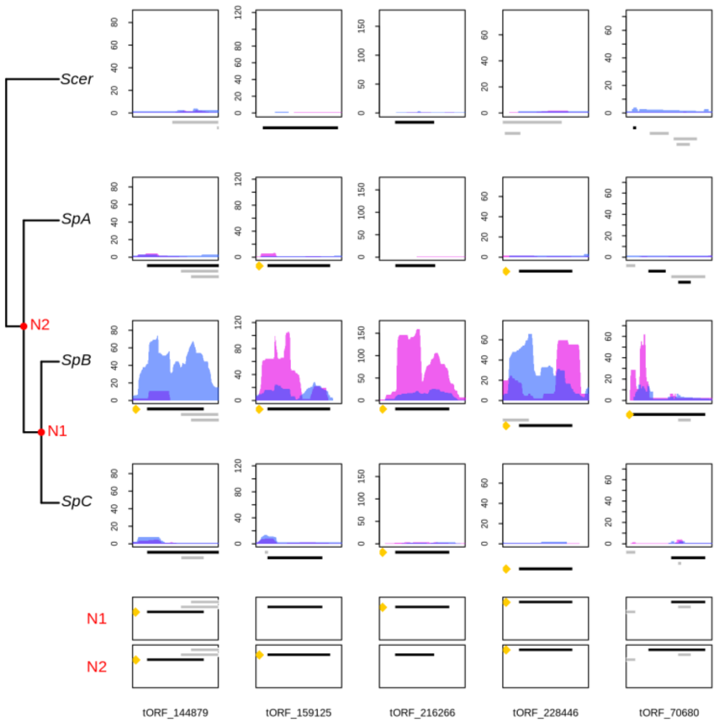


**Figure S6. Normalized RPF read coverage for lineage specific (or group specific) tORFs.** RPF read coverage is displayed for replicate 1 and 2 with a blue or pink area respectively. The positions of all iORFs (including ntORFs and tORFs) in the genomic area are drawn below each plot. The tORF of interest is labeled with a yellow dot and is plotted in black. iORFs overlapping the iORF of interest are plotted in black when they are in the same reading frame, and in grey when they are in a different reading frame as the selected tORF.

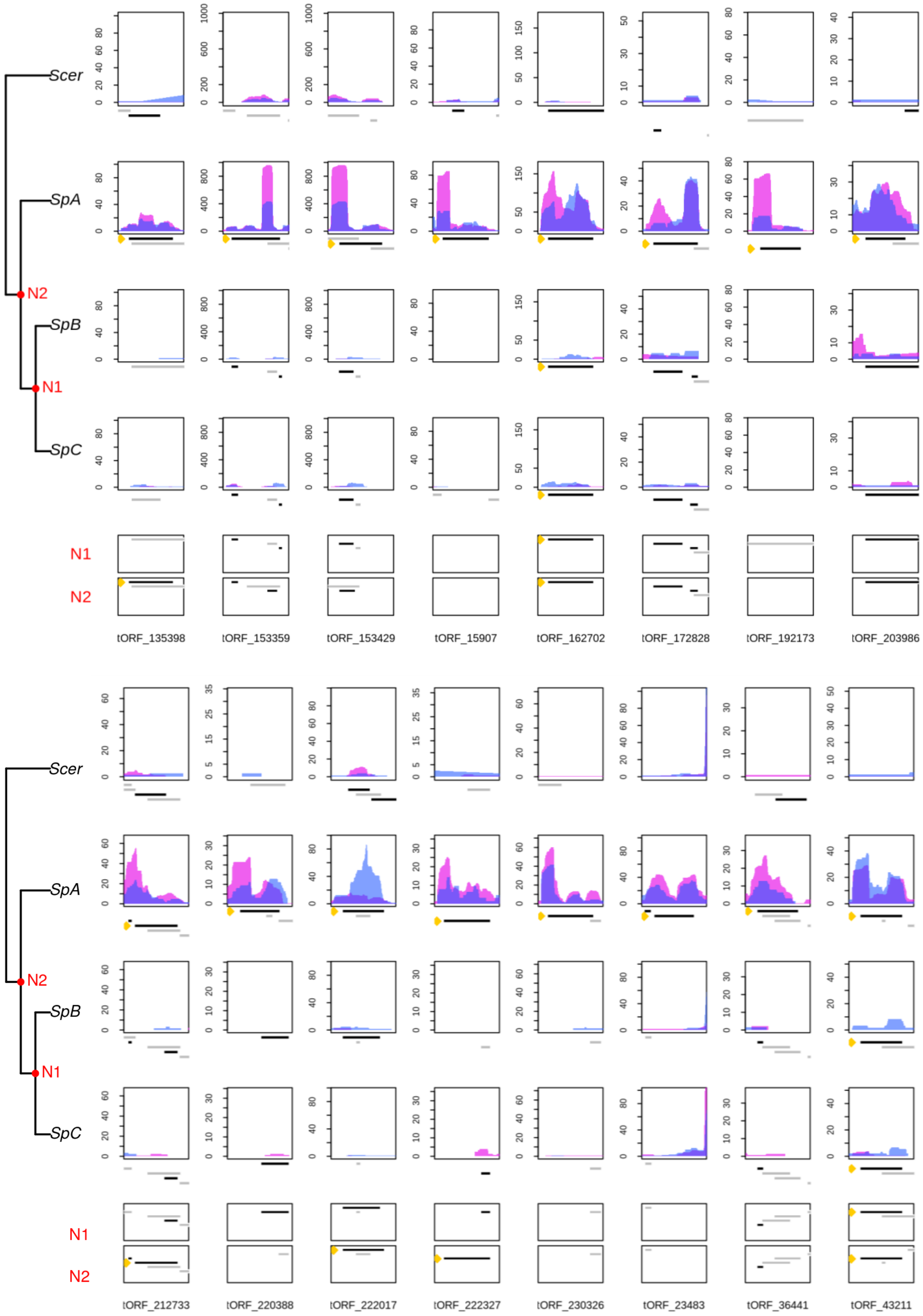


**Figure S6. Continued.**

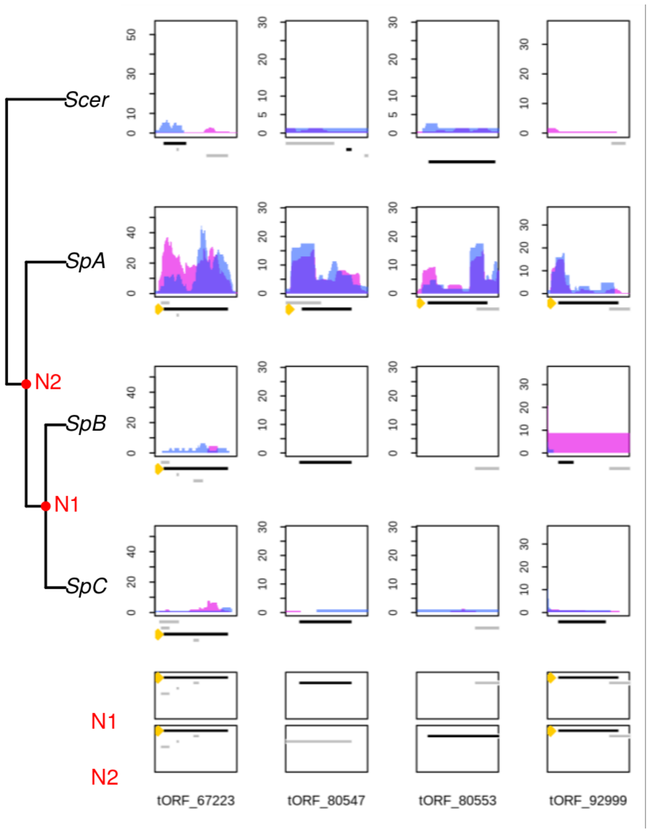


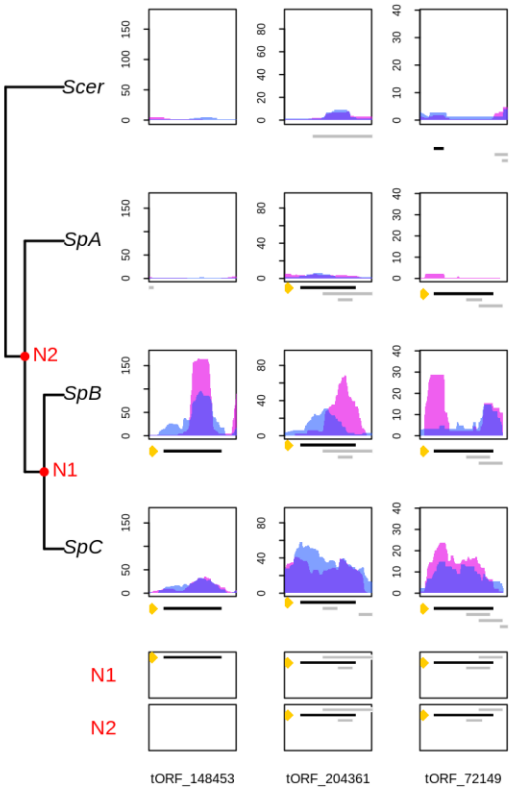


**Figure S6. Continued.**

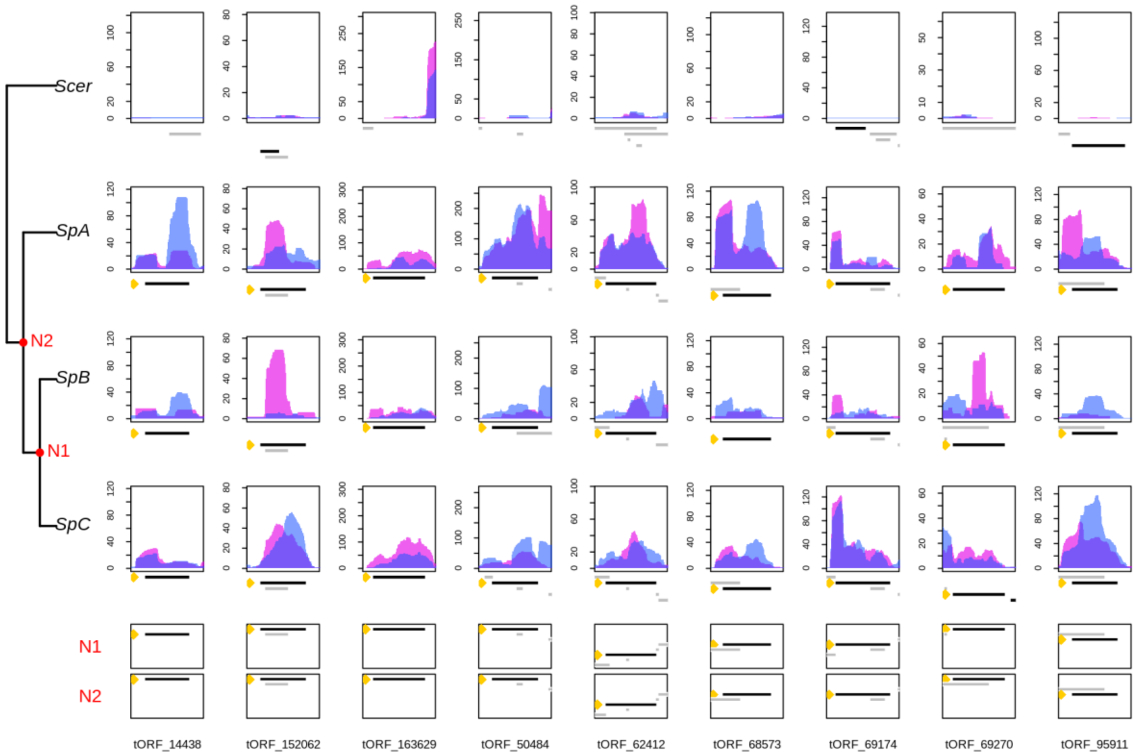


**Figure S6. Continued.**

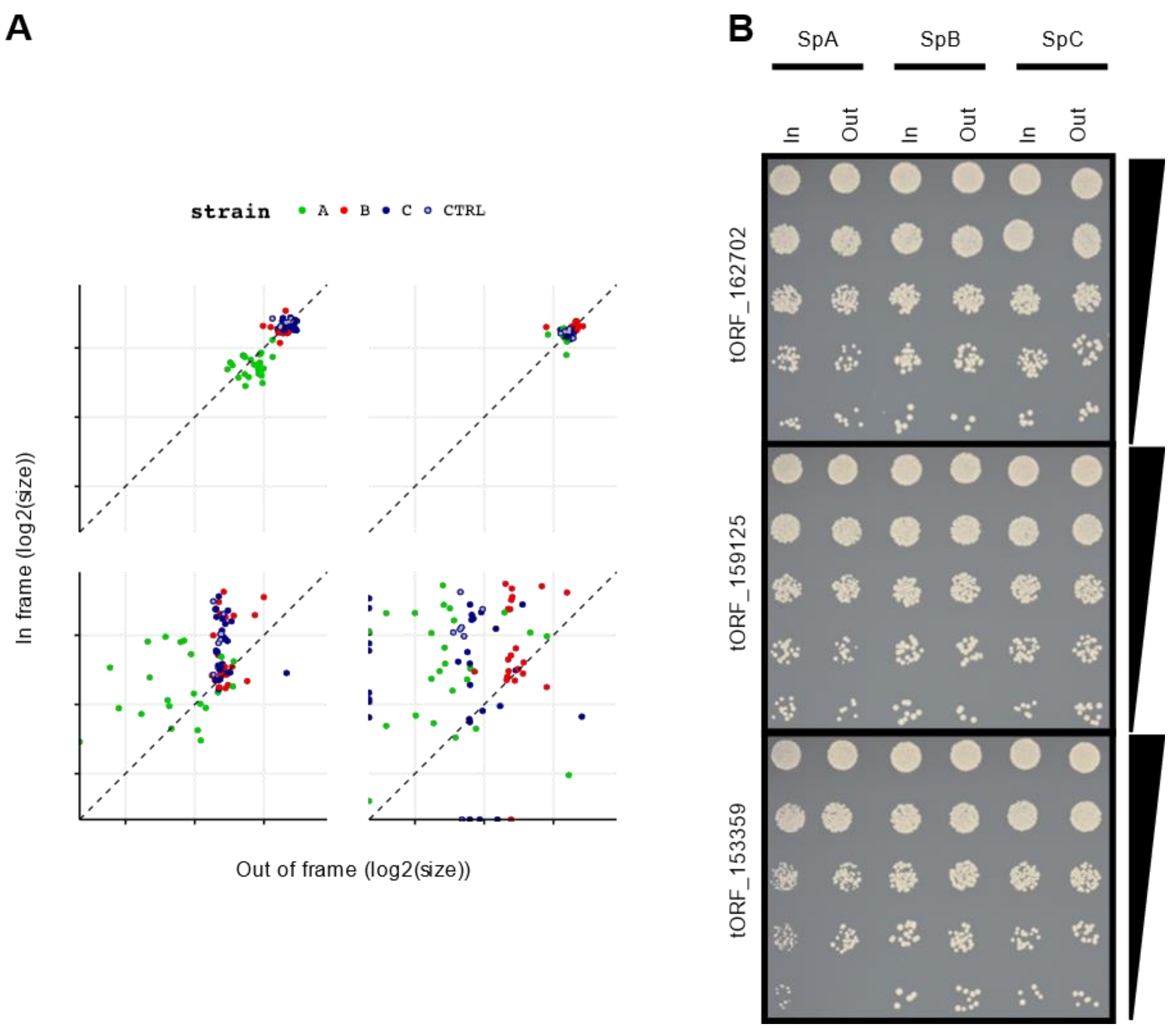


**Figure S7. DHFR tagging confirms the translation of tORFs (control experiments).** This figure is related to Fig. 6 in the main text. **A)** as in Fig. 6 B, it shows the $log_2$ colony size of strains tagged either in (y-axis) or out of frame (x-axis) with DHFR that confers resistance to methotrexate (MTX). Here, results are shown with (MTX) and without (con) methotrexate added to the media, and the data for the first (1) and second (2) phenotyping run. **B)** The control experiment for the spot dilution assays in Fig. 6C. These are the same strains, spotted at the same time from the same dilutions as in Fig. 6C but in media not supplemented with methotrexate. It shows that strains do not have growth defects.

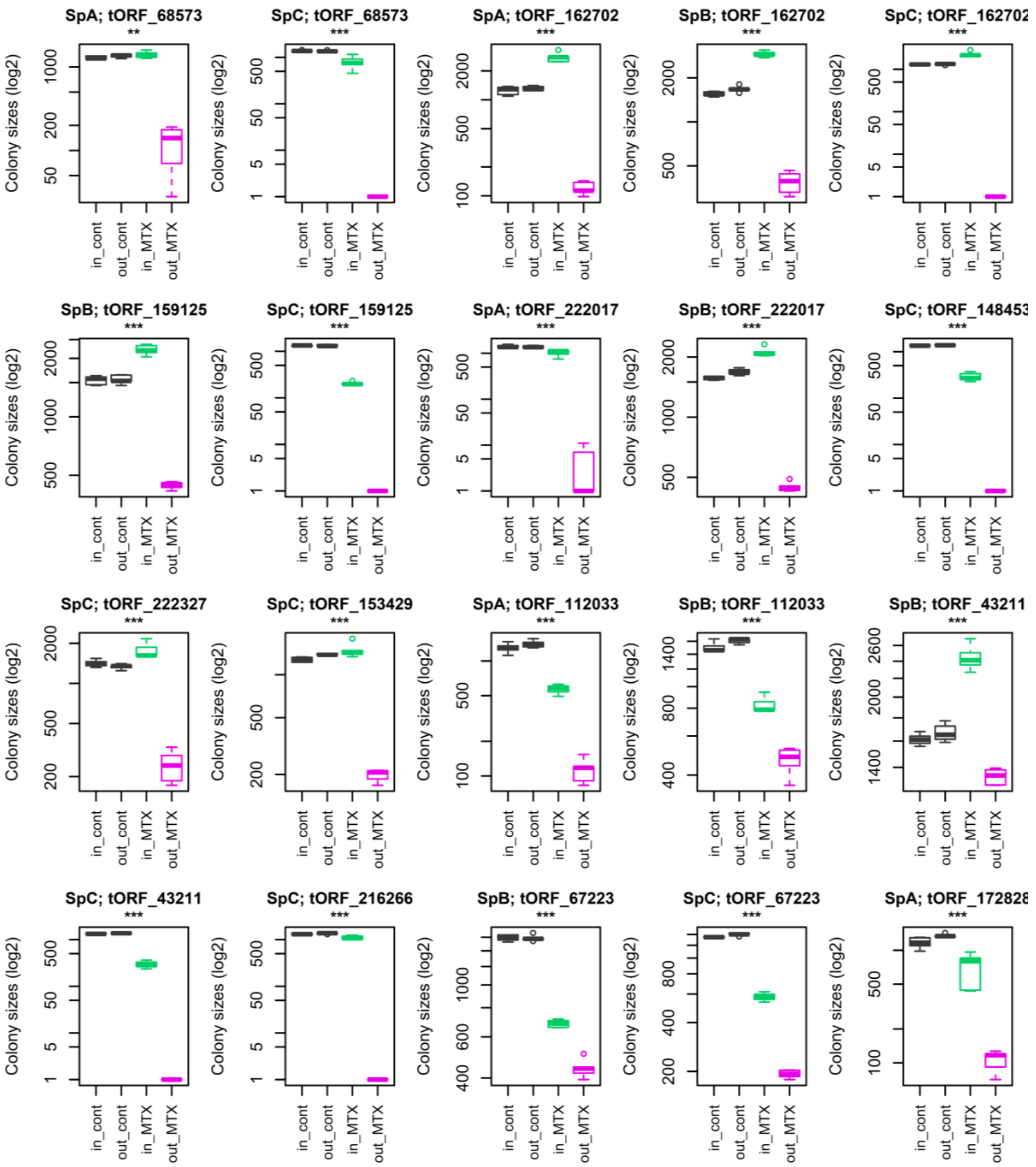


**Figure S8. Detection of translated tORFs using the reporter gene DHFR.** Each plot displays growth assay results per tORF and lineage (see headers). Note that we did not always get tORFs constructions in the three lineages so some tORFs were tested only in one or two strains (Supplemental Table S5). Positive controls (in frame and out of frame without MTX medium) are in grey, negative controls (out of frame in MTX) are in pink, and the tORFs tested for translation (in frame in MTX) are in green. 6 replicates were performed in each case. Translation was detected i) when we observed $\log_2$ colony size differences between in frame and out of frame constructions on MTX medium with a student t-test, and ii) if both positive controls display colony sizes of more than 1000 and with similar growth for both controls. Significant translated tORFs are indicated by *** for p-values < 0.001, ** for p-values < 0.01 and * for p-values < 0.05. Significant differences indicated between parenthesis correspond to either tORFs with colony size differences between controls, or to a significant higher colony size on MTX for the out of frame construct, compared with the in frame. Both situations were not considered as a translation signal for the tORF tested, the latest situation is probably due to a translated overlapping iORF.

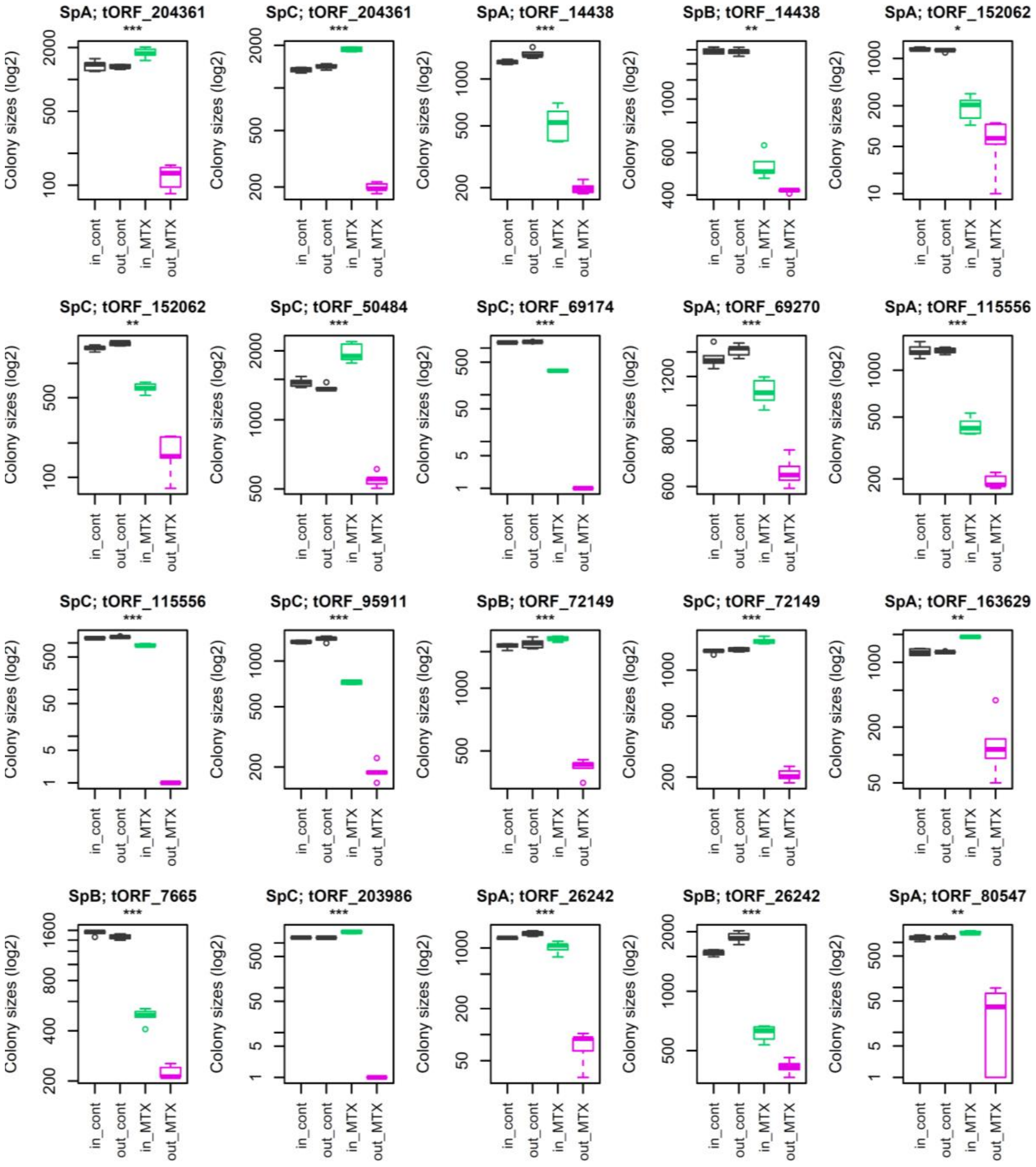


**Figure S8. Continued. Detection of translated tORFs using the reporter gene DHFR.**

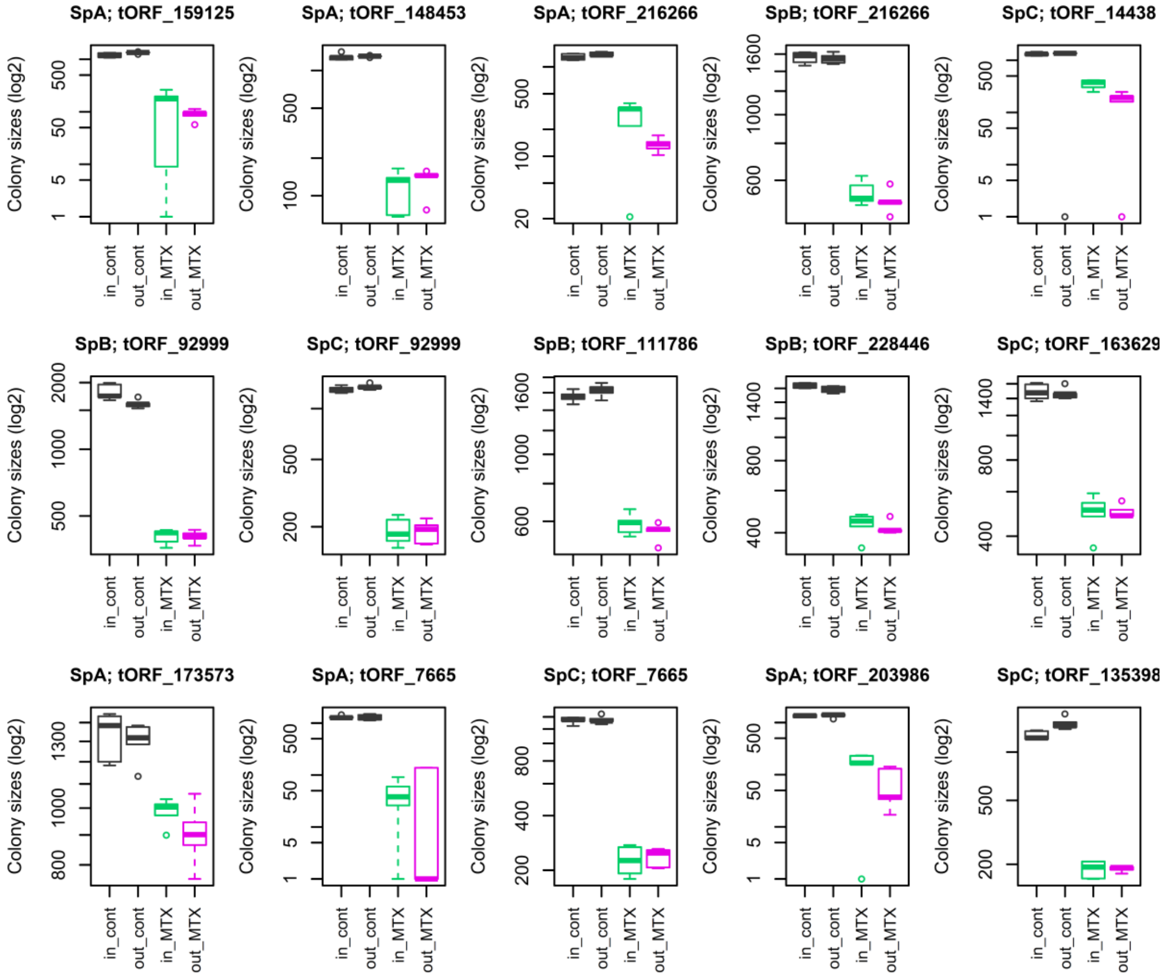


**Figure S8. Continued. Detection of translated tORFs using the reporter gene DHFR.**

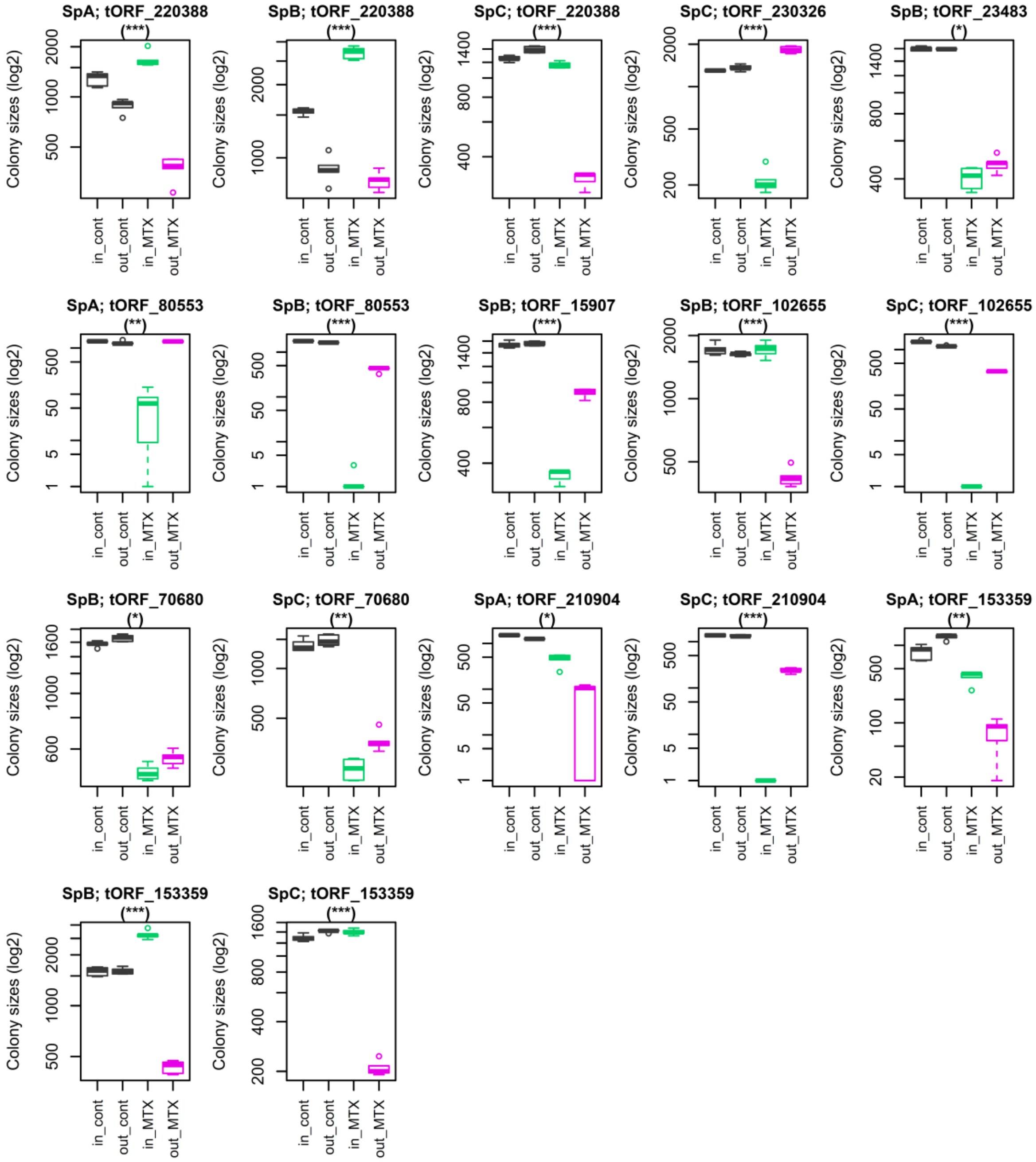


**Figure S8. Continued. Detection of translated tORFs using the reporter gene DHFR.**

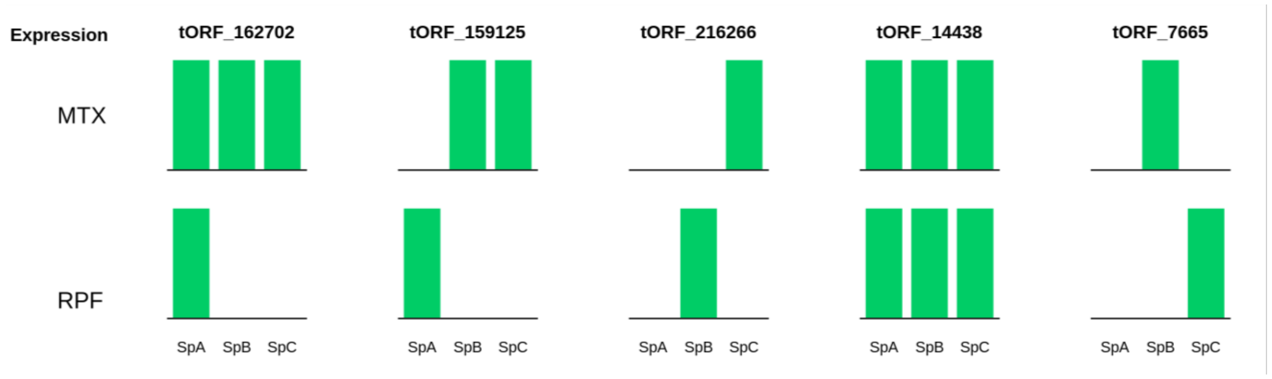


**Figure S9. Comparison of translation signals detected with the growth assay (MTX, above) or by RPF sequencing (below) in the three *S. paradoxus* lineages.** Each bar represents the detection of a translation signal per strain and per tORF, no bar is an absence. Here, we see that some tORFs are translated only under specific conditions (i.e. tORF_216266), others are more broadly translated (i.e. tORF_14438). Growth conditions therefore appear to have a strong effect on the expression of tORFs. In addition, the DHFR fusion approach could be more sensitive that the RPF approach for detecting translation.

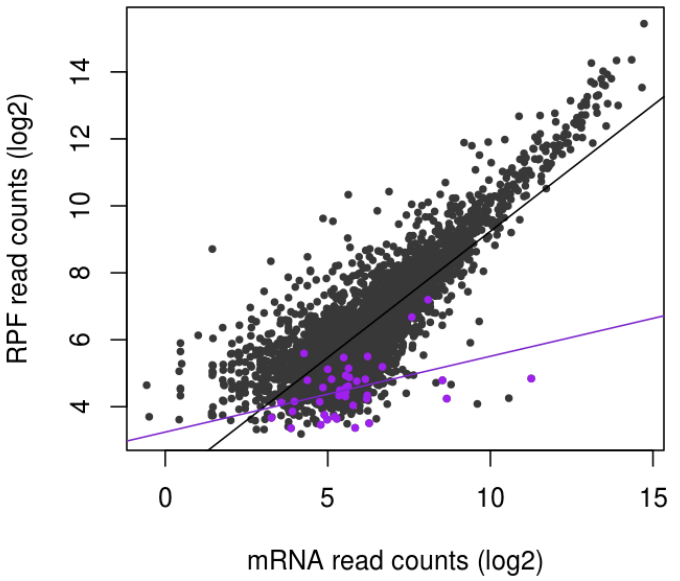


**Figure S10. TE buffering in *S. cerevisiae*.** Read counts from ribosome profiling (RPF) plotted as a function of read counts total RNA for total RNA for tORFs in purple, or genes in dark grey using ribosome profiling and mRNA sequencing from (McManus et al. 2014). Read counts were normalized to correct for library size differences. tORFs were identified based on iORFs annotations on the S228C reference strain (including *Scer* specific annotations), using the same procedure as in our analyses. We detected 40 tORFs in this dataset, which is smaller compared to our data probably due to a lower RPF coverage here. Regression lines are plotted for significant Spearman correlations (p-values < 0.05).

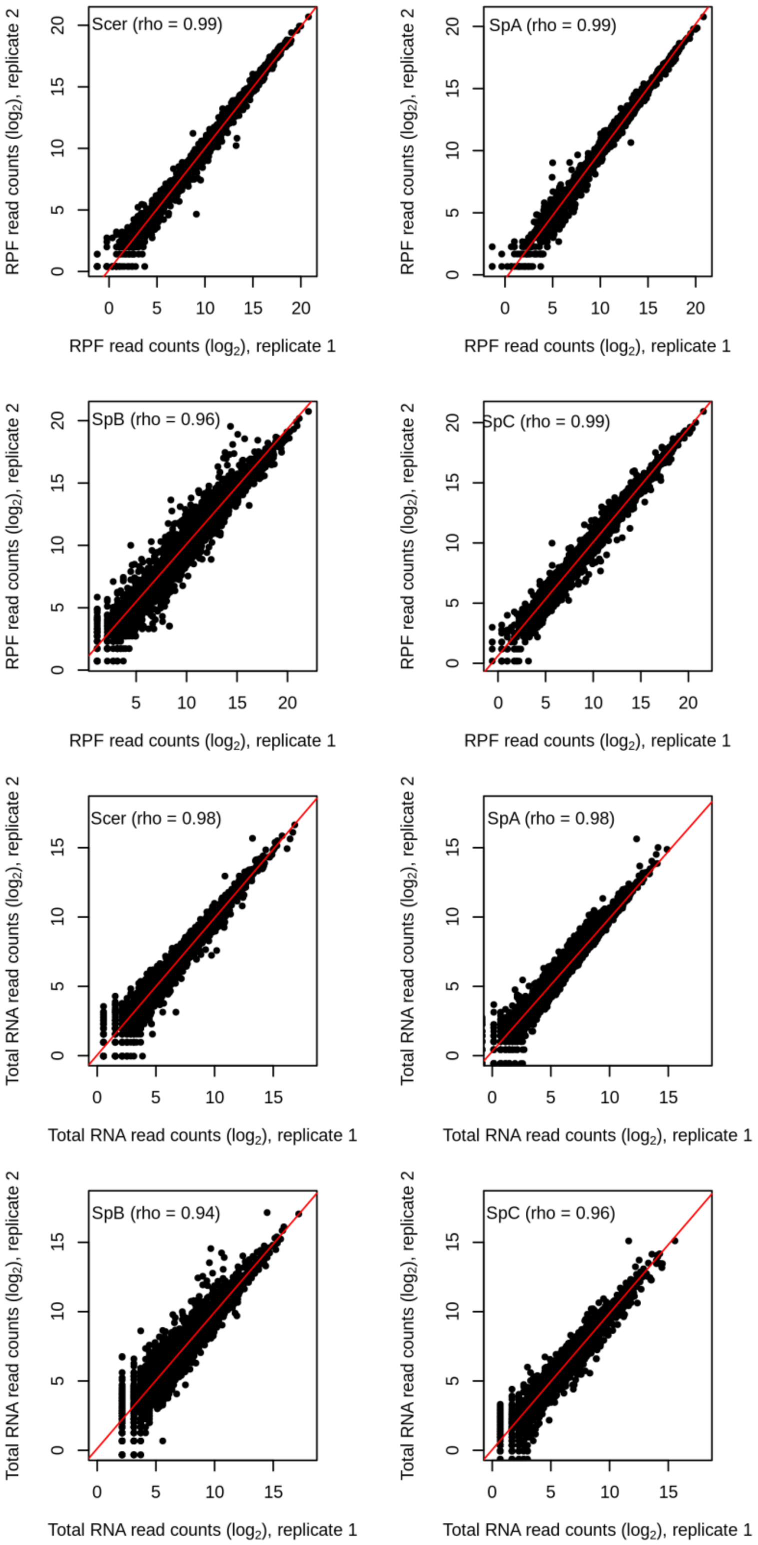


**Figure S11. Comparison of RPF or Total RNA replicate sequencing experiments.** The number of normalized read counts for biological replicate 1 is plotted against biological replicate 2 for each gene or tORF, per library type (RPF or Total RNA) and per strain. The Spearman's rank correlation coefficient (rho) is shown for each plot.

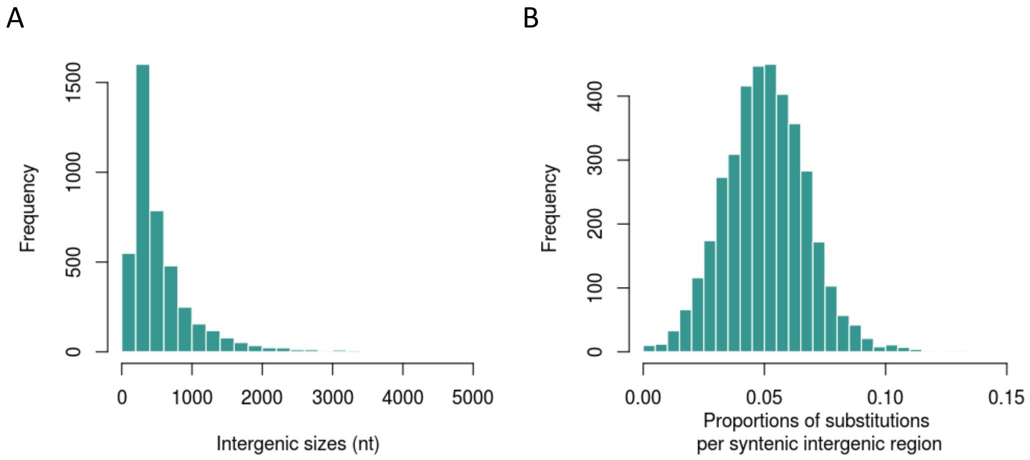


**Figure S12. Sequence characteristics of intergenic regions used for iORF annotation. A)** Distribution of the size of intergenic sequences used for iORF annotation. **B)** Distribution of SNP density per intergenic sequence.

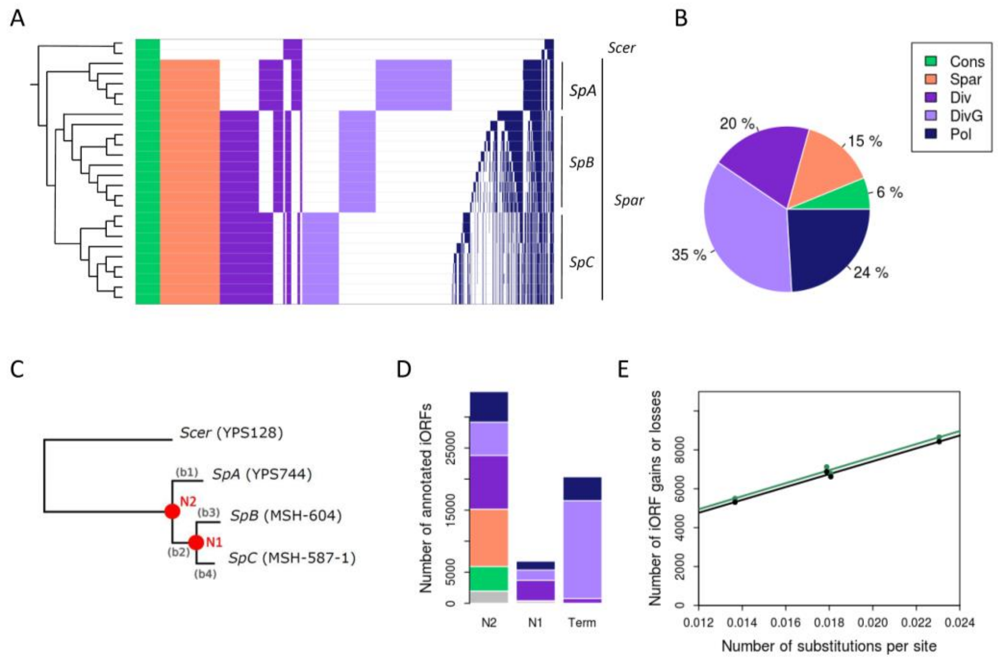


**Figure S13. Evolution of iORFs in *Saccharomyces paradoxus* populations. A)** Columns represent iORFs sorted according to their conservation levels. iORFs that are absent are shown in white. Others are colored according to their conservation group (see Methods and Fig. S1): conserved (cons), *S. paradoxus* (Spar) specific and fixed, divergent (Div), divergent group-specific (DivG) and polymorphic (Pol). **B)** Percentage of iORFs belonging to each conservation group. **C)** Phylogenetic tree of strains used for the reconstruction of ancestral intergenic sequences. Node and branch names are indicated in orange and grey respectively. **D)** Number of annotated iORFs per age, corresponding to oldest node in which they were detected. 'Term' refers to iORFs appearing on terminal branches and absent in ancestral reconstructions. iORFs detected only in ancestral sequences are plotted in gray. **E)** Number of iORF gains (in green) or losses (in black) as a function of the number of substitutions per site. Points show iORF counts on each phylogenetic branch (b1 to b4) in our dataset.

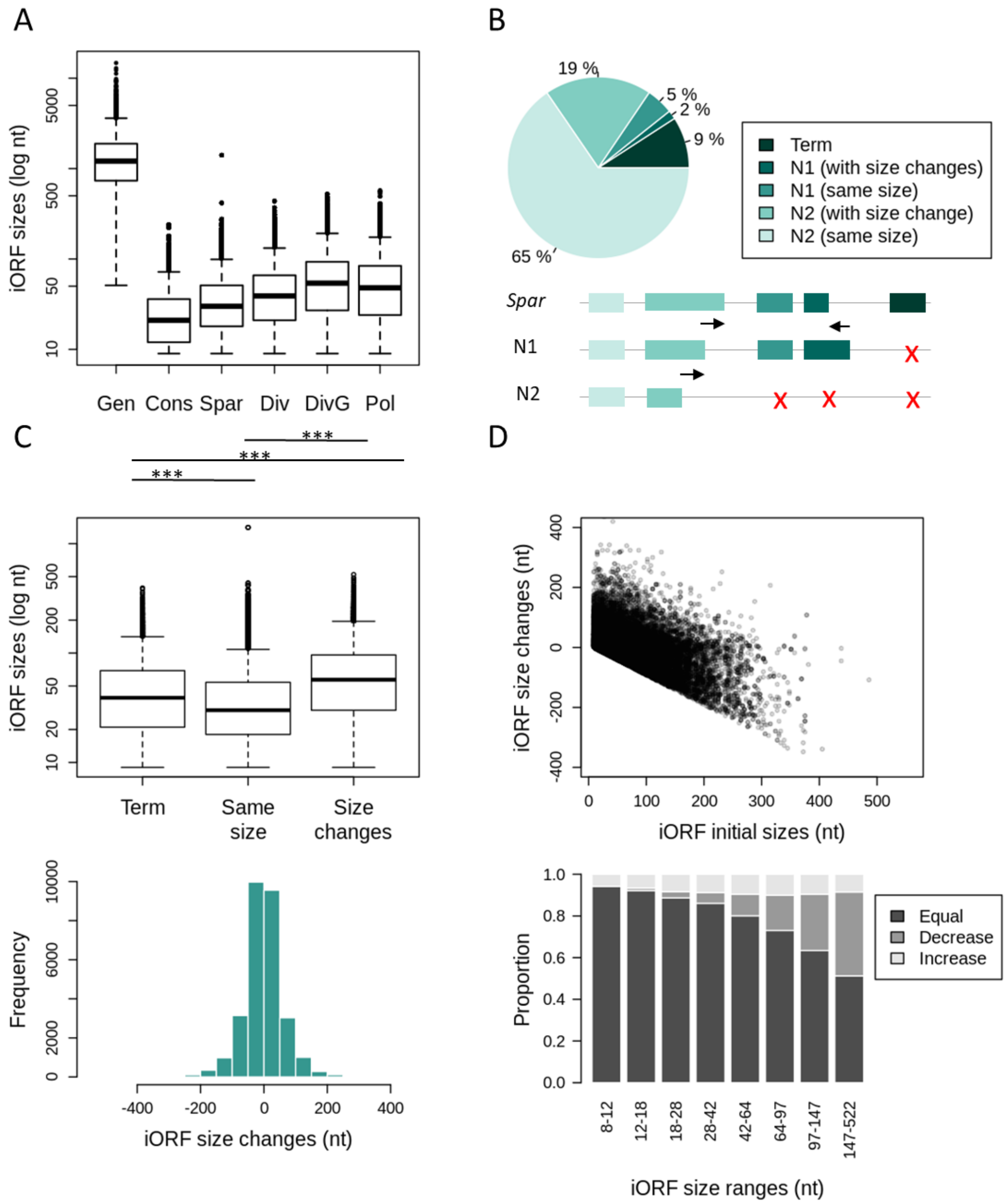


**Figure S14. iORFs change their coding potential by frequent size changes. A)** Genes (from the reference genome of *S. cerevisiae*) and iORF sizes (in nucleotides and log scale). **B)** Proportion of iORFs identified in *S. paradoxus* extent sequences (*Spar*) and successively connected to ancestral nodes, or with a connection stopped in N1, or without connection in ancestral sequences (Term). **C)** Distributions of iORF sizes in extent strains depending on the connection type with ancestral homologs: 'Term' refers to iORFs appearing along terminal branches and with no connection in any ancestral homologs, 'Same size' refers to iORFs present in ancestral sequences with no size changes and 'Size changes' refers to iORFs connected to the ancestral sequence by its start or stop position, and submitted to at least one size change. Significant pairwise differences are indicated above each comparison (Wilcoxon t-test, *** for p-values < 0.001). **D)** Size changes orientation and amplitude depending on the initial iORF size at N2 or N1. ORFs that did not change size were not displayed for more clarity. **E)** Distribution of iORF size changes relative to ancestral iORFs, and excluding ~ 16,000 iORFs, with no size changes to focus on the amplitude of changes. **F)** Proportion of terminal iORFs submitted (or not) to size changes (increase or decrease) per size range relative to their ancestor at N1.

#### Supplementary tables

Supplemental_Table_S1.txt iORF diversity in wild yeast populations.

Supplemental_Table_S2.txt iORF and gene properties (iORFs ≥ 60 nt).

Supplemental_Table_S3.txt iORF and gene raw read counts.

Supplemental_Table_S4.txt Primer DHFR tags.

Supplemental_Table_S5.txt List of DHFR constructions per strain.

Supplemental_Table_S6.txt Cell growth assay (using DHFR reporter gene to confirm *in vivo* translation).
